# Tolerant mothers: aggression does not explain solitary living in the bush Karoo rat

**DOI:** 10.1101/2024.06.24.600450

**Authors:** L. Makuya, N. Pillay, S.P. Sangweni, C. Schradin

## Abstract

Many mammal species are thought to adopt solitary living due to mothers becoming intolerant of their adult offspring as well as social intolerance between adults. However, field studies on how solitary mammals interact are rare. Here we show that solitary living can occur without social intolerance. Over three years, we recorded interactions between free living bush Karoo rats (*Otomys unisulcatus* ) and conducted dyadic encounter experiments between kin and non-kin female neighbours, both in a neutral test arena and in field intruder experiments. Social interactions were rare (N=230 in N=2062 observations), and, when observed, were aggressive in only 34% of the cases. In dyadic encounters, mothers interacted amicably with young offspring. Aggression between mothers and offspring was almost absent. This mother-offspring relationship remained amicable even after the offspring had dispersed. Aggression between neighbouring adult females was low in neutral arena tests, independent of kinship and season. However, in the field, females reacted more aggressively towards non-kin than kin intruders, especially during the breeding season. Tolerance between mothers and adult offspring indicates that aggression is not the mechanism leading to natal dispersal and solitary living. We found a solitary social system characterised by social tolerance, suggesting that dispersal in combination with a lack of social attraction rather than aggression can lead to solitary living.

## Introduction

To understand the diversity of social systems, many studies have focused on pair and group- living species, assuming solitary living to be the ancestral stage that does not require any explanation for its occurrence [1–3]. Solitary, pair- and group-living refer to the social organisation, describing the composition of social units [4]. Social organisation is one of four components of the social system which further includes the care system, mating system and social structure [4, 5]. Of all the different forms of social organisation, solitary living is the most understudied [2].

Previous studies regarded solitary living as the ancestral default form of social organisation in mammals [1]. However, recent comparative studies have shown that this is often not the case [6, 7]. These studies showed that pair-living was most likely the ancestral form of social organisation in artiodactyls [8], primates [9], and possibly in marsupials [10] and Eulipotyphla [11], indicating that solitary living is often a derived state [12]. However, we know little about the mechanisms leading to solitary living in mammals [13]. To understand the mechanisms of group-living, we also must understand the alternative, which is solitary living [14].

It is generally assumed that solitary living in mammals is due to aggression. This opinion is based on work done on the European hamster (*Cricetus cricetus*) in the 1950s [15]. In this species, mothers become intolerant of their offspring when they reach puberty and adults are highly intolerant of each other [15]. The assumption that social intolerance is the main reason of solitary living in mammals was further based on standardized laboratory experiments on solitary rodents to examine the proximate mechanisms of aggression [16–18]. However, whether this is always the mechanism leading to solitary living under natural conditions is unknown. One of the main reasons for solitary living is high reproductive competition between females which often leads to female infanticide [19–21]. As an alternative to social intolerance and aggression, a lack of social attraction combined with a motivation to disperse when reaching sexual maturity could lead to a solitary lifestyle. In solitary mustelids, for example, individuals commonly meet in a non-aggressive context [22]. Social interactions in nature have been studied for a few solitary living species such as the puma (*Puma concolor*) [23] and the giant Kangaroo rat (*Dipomys ingens*) [24]. While social interactions in these solitary species were rare, these studies highlight the lack of aggressive interactions when individuals met. More field studies are needed to understand the mechanisms that cause solitary living.

Solitary species can have complex social structures in which individuals interact in a non- random way [13]. They often display kinship-determined spatial patterns where kin live close to each other and share part of their range and engage in non-aggressive interactions when they meet [23, 24]. The kinship patterns in these solitary species are driven by philopatry, a behaviour usually displayed by group living species, with individuals dispersing only short distances and forming kin clusters [25]. For example, the social structure of the giant kangaroo rat is formed by female kin neighbours, with shorter distances between neighbours leading to increased social interactions [24]. The frequency of social interactions was positively related to population density, indicating an influence of season on the social and spatial structure [26]. Amicable interactions at territory boundaries are not a contradiction to the theoretical assumption that aggression is a main driver of solitary living, but could simply be due to the dear enemy phenomenon, where individuals are tolerant of known neighbours as long as they do not cross territory boundaries [27]. To test whether solitary living can arise without aggression, one would need to measure aggression also inside territories and at nesting sites. Field experiments are needed to test the level of aggression in solitary species, and whether close kin are more tolerant towards each other than non-kin.

Our aim was to test the general assumption that aggression and social intolerance are the mechanisms leading to solitary living in our study species, the bush Karoo rat (*Otomys unisulcatus*) from South Africa. In particular, if aggression is the main mechanism leading to natal dispersal and solitary living, we predicted (1) that the behaviour of mothers changes towards their offspring as the offspring become older, with mothers showing more aggression towards dispersed adult offspring than juvenile offspring still living with the mother. Since our study species has a kin based spatial structure with significant overlap of home ranges between close kin [28], we further predicted (2) that they would show higher levels of aggression towards non-kin neighbours than towards kin neighbours. To test these predictions, we conducted more than 2000 focal animal observations over a period of 3 years, conducting tests in a neutral presentation arena in a field laboratory, and conducted field experiments where we presented kin and non-kin neighbours at the nesting sites of resident females.

### Materials and methods Study site

The study was conducted in the Goegap Nature Reserve, in the Northern Cape, South Africa. The study site is located in the Succulent Karoo biome [29]. The climate is arid with temperatures falling below 0°C in winter and exceeding 40°C in summer [30]. Mean precipitation at the field site is 160mm per annum. Seasons are divided into the hot dry non-breeding season (December to May) and the cold wet breeding season (June to November). The bush Karoo rat reproduces in the wet season.

### Study species

The 100g heavy bush Karoo rat offers a model to study solitary living because it is diurnal, occupies an open habitat, has small home ranges (0.06 ± 0.04ha in the dry, non-breeding season and 0.04 ± 0.03ha in the wet, breeding season) [28], and easily habituates to the presence of observers. It inhabits the semi-arid regions of South Africa including the Succulent Karoo (less than 200mm of rain per annum), one of the world’s most important biodiversity hotspots [31]. The bush Karoo rat builds stick-lodges inside shrubs, which offers a favorable microclimate with high humidity and mild temperatures as protection against the harsh outside environment that is characterized by unpredictable rain in cold winters, and long, hot summer dry seasons [32–34]. The bush Karoo rat is a central place forager, foraging around its stick lodge and bringing food back to the lodge, where it can be easily observed, and where experiments can be conducted. In the Succulent Karoo, around 95% of the rats are solitary, although a few small groups of 2-3 closely related females occur [28]. They have a kin based spatial structure, with females having more kin than non-kin neighbours, and home ranges of close kin overlapping more with each other than of non-kin [28]. Young male bush Karoo rats behave similarly to females at the beginning of the dry season and stay in an area close to their natal lodge. However, they disperse in winter when food availability increases just before the breeding season starts. Adult males roam over very large areas and there are no resident males on the field site during the breeding season. Thus, there are no male neighbours in the breeding season.

## Sampling regime

### Marking and trapping

Trapping was conducted at lodges that showed signs of being occupied (fresh faeces, active runways, rats observed). The field site was divided into six areas, with 1-2 areas trapped simultaneously by research teams. All lodges within one area were trapped for three consecutive days before moving on to the next area. Traps were set in the morning before sunrise and checked after 45 and 90 minutes and then un-set during the hottest times of the day. In the afternoon, traps were set 45 minutes before sundown, checked after sundown, and then un-set during the night. We used a combination of foldable Sherman traps (https://shermantraps.com/) and locally produced heavy metal traps of the Sherman style. Traps were baited with a combination of bran flakes, salt and sunflower oil and re-baited each morning and afternoon. Traps were arranged around the entrances of the lodges and along runways. We recorded the body weight of individuals to the nearest 0.1-gram, as well as their sex, reproductive status and lodge number. We marked individuals with single, metal-band ear tags with a unique reference number (National Band and Tag Co., Newport, KY, U.S.A.) [35]. To aid in visual identification during observations, individuals were marked with non-toxic hair dye (Inecto Rapido, Pinetown, South Africa), in combinations (females: head and chest/sides/back; males: hindquarters and chest/sides/back). Age was estimated from body mass at first capture, using a species-specific growth curve [36] validated for our field data. The bush Karoo rats were classified according to their age, with pups being up to two weeks old and weighing less than 30 g (weaning is at 14 days; [36]), juveniles being 2-6 weeks old and weighing between 30 and 70 g, and adults being older than 6 weeks and weighing more than 70 g, when both sexes can start reproduction [36].

### Determining dispersal

The onset of dispersal was determined for the rats used during the dyadic encounter tests (explained below). We determined these occurrences from the time that the rat was trapped consistently over a period of more than 4 weeks at a lodge that was not its natal lodge, without the mother being trapped and observed at the same lodge. Using these data, we calculated the age at dispersal.

### Focal animal observations in the field

Focal animal observations were conducted between March 2021 and October 2023 for a total of 2279 observations for 246 rats at lodges with identified bush Karoo rats to establish which individuals occupy the lodges, and 2) record behaviours, including interactions with conspecifics. The observations were done for 30 minutes in the morning after sunrise and 30 minutes in the afternoon before sunset. Observations were done using focal animal sampling and one zero recording for 30min. We recorded all social behaviours in 1-minute intervals. The social behaviours including the following groups of behaviour: (i) amicable behaviours (e.g., grooming, body contact); (ii) social investigation (i.e., sniffing); and (iii) aggression (e.g., chasing, fighting).

### Dyadic encounter tests

Dyadic encounter experiments were used to assess whether interactions with neighbours were amicable (predicted for close kin) or aggressive (predicted for non-kin). Bush Karoo rats were trapped and brought inside their trap to a laboratory at the research station, located 100 m away, and allowed to acclimatize for 10 minutes before the start of the experiment. The experiments were conducted in a neutral test arena that was constructed of wood chip panels (80 cm x 65 cm x 94 cm) and had a partition in the middle (Figure S 1, supplementary file). The testing arena was cleaned between encounters using diluted Dettol Antiseptic Liquid and then air dried. All tests were done between 10 am and 12 pm from August 2021 until March 2023, for a total of 143 tests on 52 focal rats. Each rat was tested an average of 2.80 ± 2.2 times.

Each bush Karoo rat was introduced into the arena and allowed to settle for 5 minutes with the partition down. We first tested mothers as the focal individual against their offspring which were between one month and 20 months old (N = 80 tests, 3 with male offspring and 77 with female offspring, on 31 focal mother rats). The offspring tested were either still living in their mothers’ lodge i.e., not dispersed (N = 43) or already dispersed (N = 36). We attempted to test each mother with the same offspring at different ages, but because some rats were not re-trapped and disappeared, some mothers were tested with different offspring at different ages. Next, we tested adult female focal rats on two different days with an adult female kin neighbour and an adult female non-kin neighbour respectively (N = 63 tests on 41 rats). Half of the rats were first tested with a kin neighbour, the other half with a non-kin neighbour. We defined a neighbour as a rat that occupied a lodge not more than 25 metres away from the focal rat. A total of 22 of the focal rats tested with a neighbour were used in the tests with offspring. Because all stimulus animals were direct neighbours, our experimental design controlled for the dear enemy phenomenon, whereby the owner of a territory responds less aggressively to a familiar neighbour than towards a stranger [27]. Presentations lasted for 15 minutes each. The focal adult individual was heavier than the stimulus female (mean weight difference 25.8g ± 20.3 *SD*), because we wanted the focal individual, which we assigned as the owner of the territory, to initiate the encounters and body mass difference was expected to have a positive influence on the initiation of aggression [37]. At the end of each test, bush Karoo rats were returned to their lodge. Focal animal sampling was used to record the frequency of social behaviours as described for focal animal observation above [38]. The behaviours were recorded for 15 minutes using a webcam. The observer was present in the same room as the animals being tested but was separated from the animals by a black curtain and monitored the behaviours live on a computer. Tests would have been immediately terminated as soon as individuals started damaging fights (biting and/or standing upright and boxing for more than 2 seconds) to avoid any injury. However, this was never necessary in our study. Interactions videos were then scored using BORIS (Behavioural Observation Research Interactive Software, [39]).

### Field intruder presentation tests

Since we observed little aggression during the neutral arena tests, we conducted field intruder tests directly at the lodge of the rats. A similar experiment had been done on group living African ice rats (*Otomys sloggetti robertsi* ) from the alpine regions of the southern African Drakensberg and Maluti mountains [40]. We expected higher levels of aggression here due to the focal rats defending their territory, which was not the case in the neutral arena in the laboratory. Again, we tested whether focal animals are more aggressive towards non-kin than kin neighbours. These tests were done 1-2 years after the dyadic encounter tests in the neutral test arena, and of the individuals tested in the field, only nine had participated as the focal (N = 9) / stimulus (N = 6) in the previous experiments. Stimulus individuals were trapped (as described above) and then transferred into a wire mesh cage (30 x 15 cm, 12 cm height) (Figure S2). This wire mesh cage allowed other bush Karoo rats to see and smell the stimulus animal in the trap, but not to reach and touch it. The trapped individual was then presented at a neighbouring rat’s lodge (i.e. the focal individual). The focal individual was not caged and its response to the trapped stimulus individual was recorded. The wire cage was positioned within an active runway 30 cm away from the lodge. Observations started when the focal animal was observed outside its lodge and lasted for 15 minutes thereafter. The maximum duration of the presentation was 45 min; thus, when the focal animal was not seen within the first 30 min, the experiment was terminated. During the 15 min of observations, we recorded aggressive behaviours of the focal animal towards the caged stimulus animal, including charging towards the cage and emitting chit sounds. We also recorded the latency until the investigation of the stimulus individual and the first aggression as well as the total time spent at the cage. Thereafter, the stimulus animal was returned to its lodge from where it was trapped. Each focal was tested, once with a kin neighbour and once with a non-kin neighbour on separate days. The tests were conducted in the early mornings between 6:00 and 9:00 from January – November 2023. We conducted a total of 54 tests for 26 focal rats with each rat being tested for an average of 2.03 ± 0.87 times. For both the dyadic and field encounter tests. a focal rat could be used as a stimulus in another experiment, and a stimulus rat could be used multiple times for different focal rats. In the dyadic tests, 15 rats were used as both focal and stimulus and 12 rats were used in the intruder tests.

### Statistical analysis

All statistical analyses were done in R [version 4.3.1; 41]. We used an ANOVA to analyse factors that influence the frequency of sniffing events, or the time spent in body contact, fighting, or grooming, and included the age of offspring, and season (breeding vs non-breeding) as factors for the dyadic encounter tests between mother and offspring (Table 1).

**Table 1:**
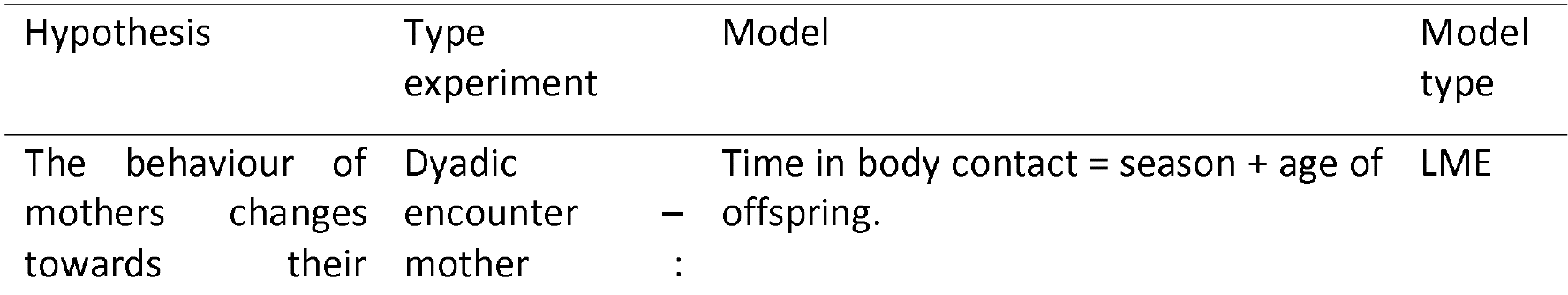
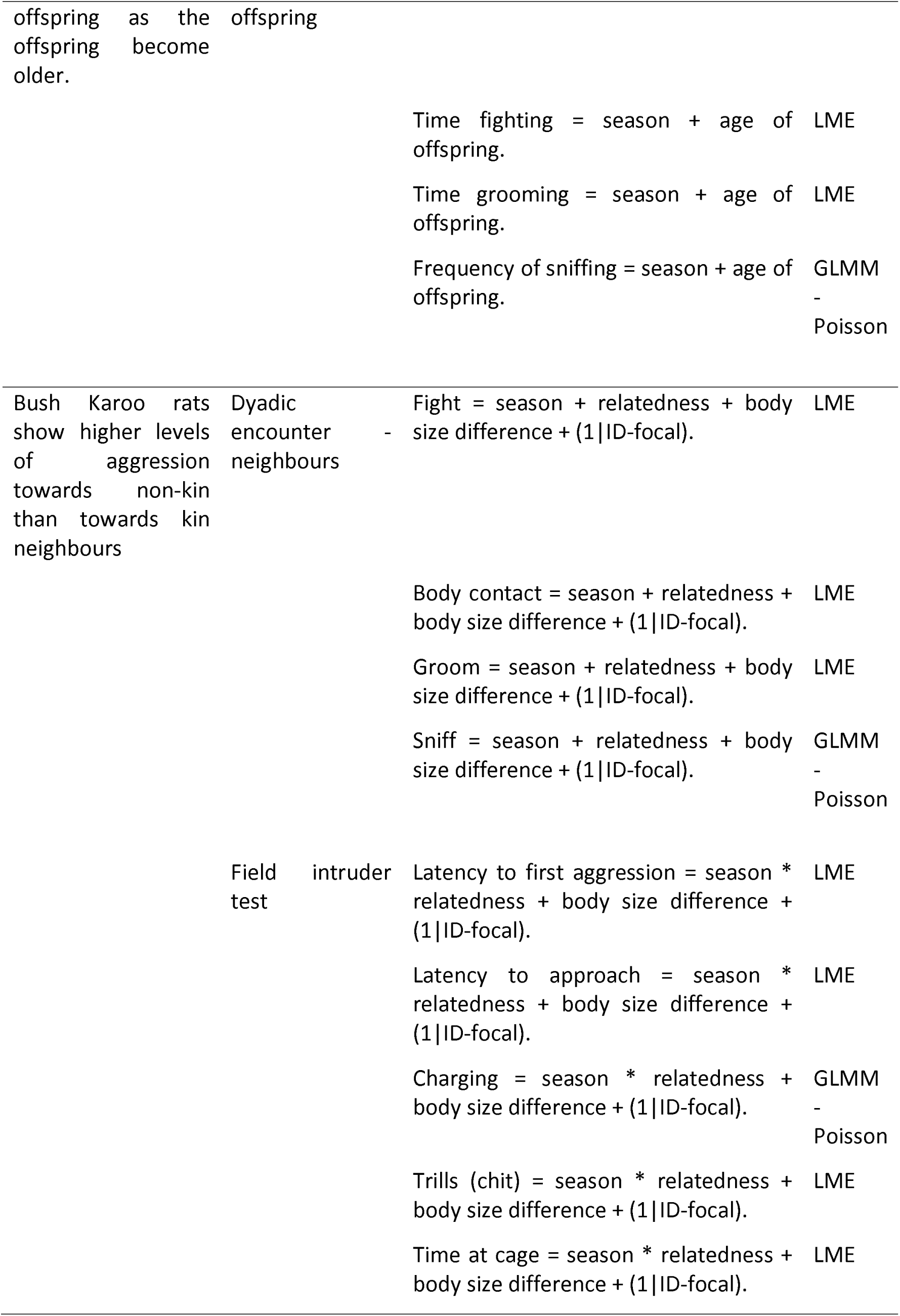
Hypothesis and associated models tested in the dyadic and field intruder tests.

We ran linear mixed models (LME) in *lme4* to investigate factors that influenced the behaviour exhibited by bush Karoo rats during dyadic encounter tests toward neighbours. For affiliative behaviours, we tested the time spent sitting in body contact, and time spent grooming. For aggressive behaviours we tested the duration of fights. Finally, for social investigative behaviours, we fit a generalized linear mixed model (GLMM) with a Poisson distribution, and we tested the number of sniffing events displayed. For all the model, the duration/frequency of the behaviours were fitted in linear models as predictors, and we included season, body size difference, and relatedness as fixed effects and the ID of the focal individual as random effects (Table 1).

We used both generalized linear mixed models (GLMM) and linear mixed models (LME) to analyse factors that influence the behaviour displayed by the focal rat towards a stimulus for the field intruder presentation tests. We tested the latency to approach the cage and to first aggression, and the number of charging events (fitted with a Poisson distribution) and the number of chit sounds, and total time spent at the cage. One model was fit for each behaviour. The behaviours were again fitted as predictors, and the interaction between season and relatedness included as fixed effects. To avoid singularity, we only fitted the ID of the focal rat as a random effect (Table 1).

## Results

### Mother- offspring interactions

During 87 of the 2062 field observations sessions, pups were present at the mother’s nest. On 11 occasions, we observed amicable interactions with the mother (grooming, and body contact), and no interactions occurred on 76 occasions; we never observed aggression towards the offspring by the mother. Juveniles were present on 240 occasions during the field observations, and were observed in amicable interactions with the mother on 20 occasions, and only once in an aggressive interaction.

The mean age of all offspring (N = 79) used as stimulus animals was 4.13 ± 3.72 *SD* months. The age of non-dispersed rats in the experiments was 3.81 ± 3.25 *SD* months (range 1 – 14, N = 43) compared to 5.21 ± 4.01 months (range: 1 – 20, N = 36) of dispersed rats. We calculated AIC values for models including the age of offspring vs dispersed or undispersed. We found that the AIC values for models including the age of offspring as a predictor were higher than for the models that included whether the offspring had dispersed (Table S2, supplementary file). However, the summary tables for both these models were very similar (results and conclusions did not change) and we reported the models including the age of offspring in the electronic supplement. Dyadic encounters between mothers and their female offspring were characterised by sniffing and body contact with little grooming and nearly no aggression (Table S1, supplementary file). Social interactions were not influenced by season and did not significantly change after offspring dispersed (Figure S3; Tables S3-S6: supplementary file). Specifically, mothers were not more aggressive towards dispersed offspring than non-dispersed (Table S4, Figure S4), nor did they decrease the level of body contact (Table S6. Figure S3).

### Focal animal observation

In 230 of the 2062 observations, conducted between July 2021 and October 2023, two or more adult rats were present at the focal rats’ lodge. Only 97 chases occurred in 79 of the 230 observations. Chases occurred more often during the non-breeding season (t-test; *t*_1728_

*= -2.*99, p < 0.01). The chases occurred between a male and an unrelated female (41%) or an unidentified neighbour (41%), and between related females (18%).

### Dyadic encounter tests (neutral test arena)

We tested whether adult female bush Karoo rats (focal individuals) showed less aggression towards kin neighbours than towards non-kin neighbours (adult females). The weight difference between the individuals was not a significant predictor in any of the models. Adult female bush Karoo rats showed very little aggression towards their neighbours, irrespective of the season (LME; estimate = 0.016, s.e. = 0.013, CI: -0.0098/0.042, *t*_63_ = 1.2, *p* = 0.22) and relatedness (LME; estimate = 0.009, s.e. = 0.015, CI: -0.02/0.04, *t*_62_ = 0.6, *p* = 0.55) (Figure S2 and Table S7: supplementary file). However, they spent more time in body contact with kin than with non-kin (LME; estimate = -2.36, s.e. = 1.316, CI: -5.04/0.31, *t*_63_ = - 1.79, p = 0.08; Figure S5 A), but the season was not a significant predictor (LME; estimate: 0.57 + s.e. = 1.094, CI: -1.63/2.8,*t*_61_ = 0.52, *p* = 0.60) (Table S8: supplementary file). The time spent grooming neighbours was significantly affected by season (LME; estimate = 0.008, s.e.= 0.019, CI: -0.0295/0.045, *t*_63_ = 0.42, *p* = 0.673) or relatedness (LME; estimate = -0.02, s.e. = 0.022, CI: -0.0645/0.025, *t*_58_ = -0.93, *p* = 0.356) (Figure S5 B and Table S9: supplementary file). Bush Karoo rats spent a significant amount of time investigating neighbours through sniffing in the breeding season (GLMM; estimate = 0.73, s.e. = 0.27, CI: 0.22/1.29, *t* = 2.697, *p* < 0.01) but this was not affected by relatedness (estimate = 0.54, s.e. = 0.36, CI: - 0.17/1.27, *p* = 0.14) (Figure S6 and Table S10: supplementary file).

### Intruder field presentation tests

In both seasons, female bush Karoo rats were mildly aggressive towards non-kin neighbours but tolerated kin neighbours when presented at their lodge. There was no difference in the latency to approach the cage with a kin or non-kin focal neighbour(estimate = -0.98, s.e. = 1.61, CI: -4.27/2.29, p = 0.6; Figure S8 and Table S12), indicating resident bush Karoo rat females investigated the stimulus animal regardless of kinship. However, non-kin females were attacked much faster (LME; *t*_38_ = 2.54, *p* <0.05, Figure 2 and Table S11), their cages were charged more often (GLMM; estimate = 0.75, s.e. = 0.22, CI: 0.32/1.2, *t* = 3.4, *p* < 0.001; Figure 2 and Table S13), and they produced more trill sounds (Figure S8; although this was not significant because a large outlier: LME; estimate = 9.53, s.e. = 18.2, CI: - 25.45/44.77, *t =* 1.56, *df* = 37.59, *p* = 0.61; Table S14). Also, focal individuals spent significantly more time at the cage with the non-kin than kin neighbours (LME; estimate = 4.63, s.e = 1.37, CI: 1.76/7.39, *t*_39_ = 4.06, *p* < 0.01, Figure. 4 and Table S15). These differences occurred in both the breeding and the non-breeding seasons. However, in the breeding season, female bush Karoo rats showed more aggression towards intruders, regardless of kinship (LME; *t* = -4.32, *p* < 0.001) by quickly attacking (LME; estimate = 4.287, s.e. = 1.82, CI: 0.61/7.95, t_43_ = 2.349, p = 0.02), and charging at cages (LME; estimate = -1.92, s.e. = 0.44, CI: -2.899/-1.10) (Figures. 2-3).

**Figure 1.**
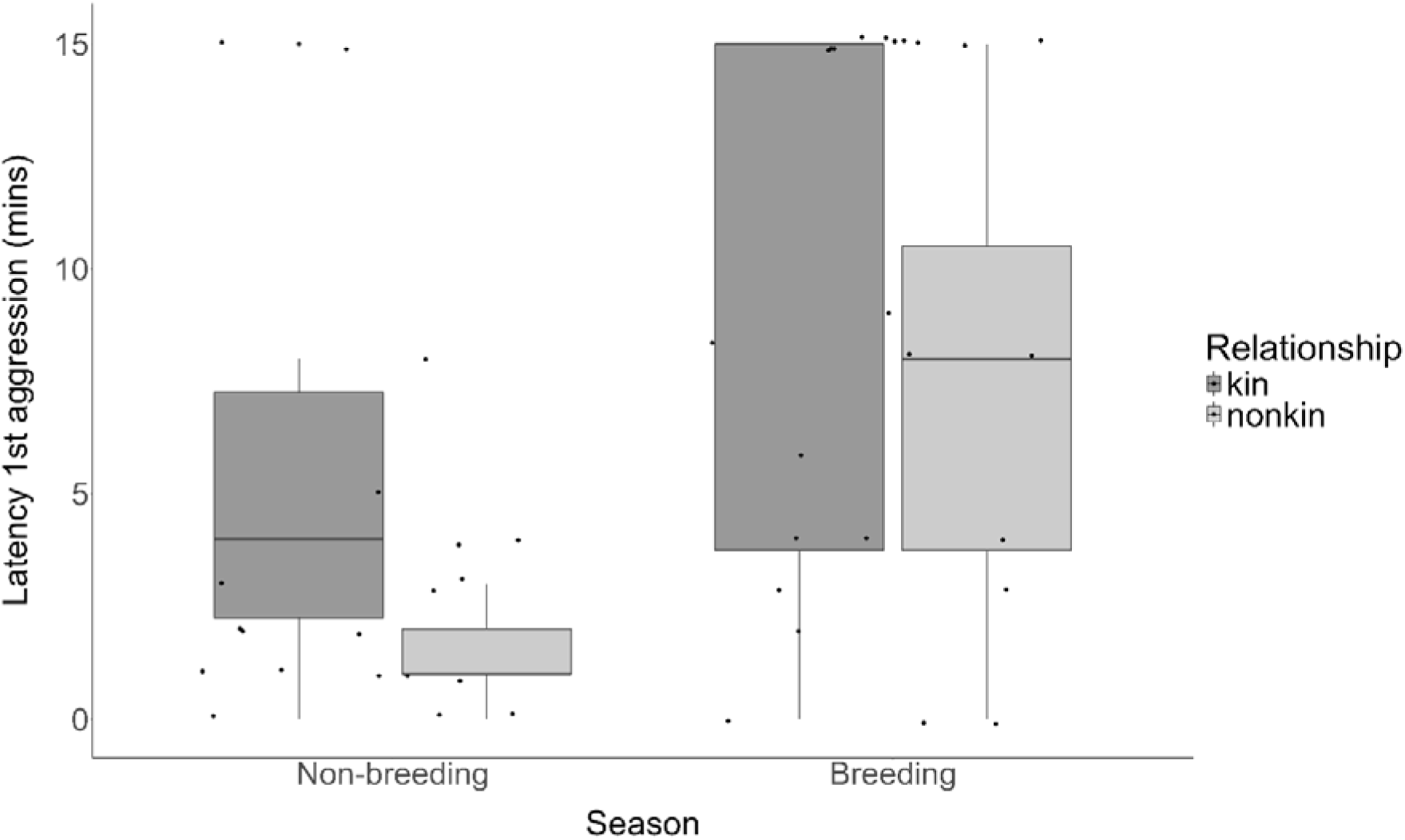
: The latency to first aggression by focal females to neighbouring cage-housed female bush Karoo rats during intruder tests. Boxplots show median and 1st and 3rd quartiles, the whiskers represent the minimum and maximum of the outlier data, and points represent individual values.

**Figure 2:**
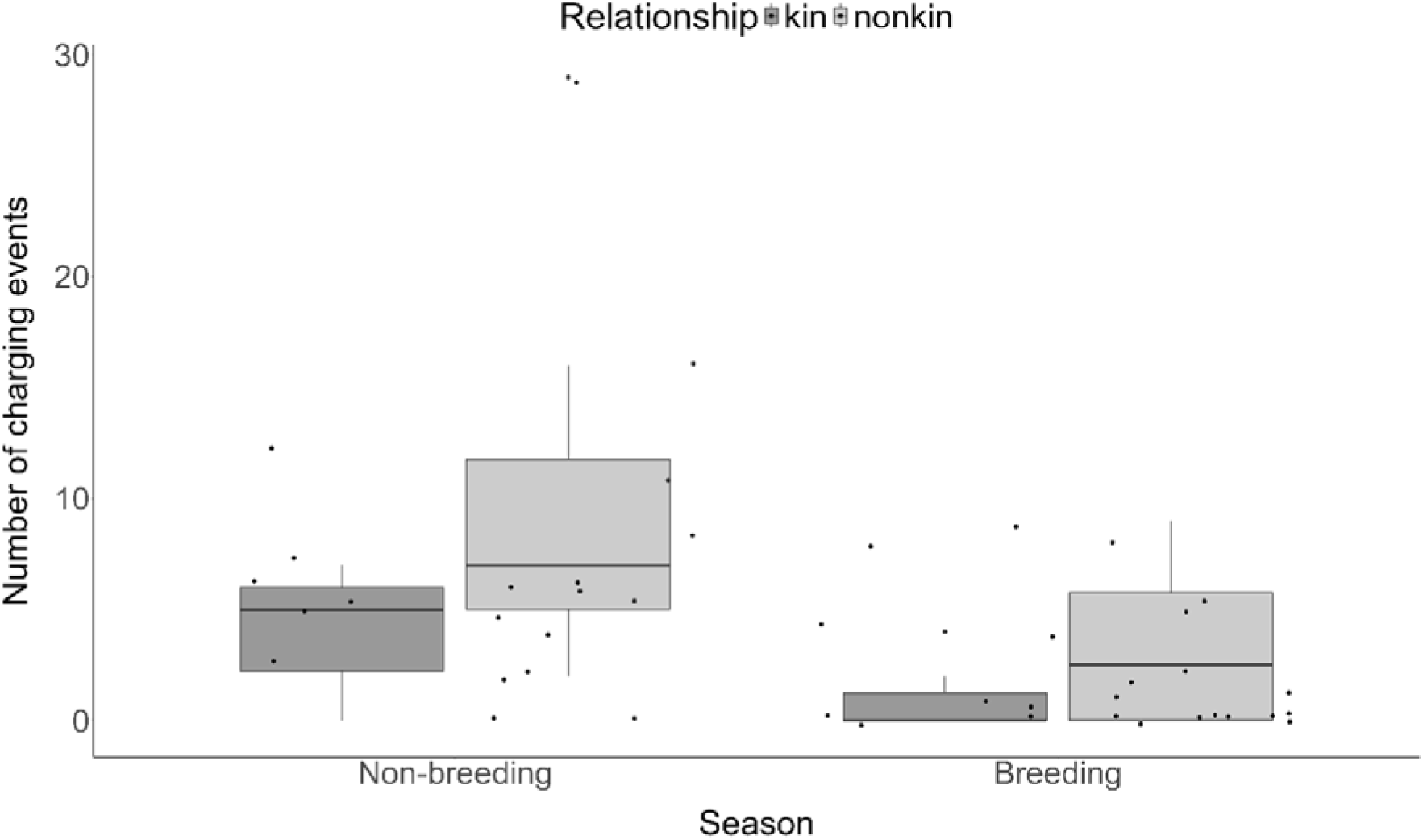
The number of charging events by focal females to neighbouring cage-housed bush Karoo rats during intruder tests. Boxplots show median and 1st and 3rd quartiles, the whiskers represent the minimum and maximum of the outlier data, and points represent individual values.

**Figure 3:**
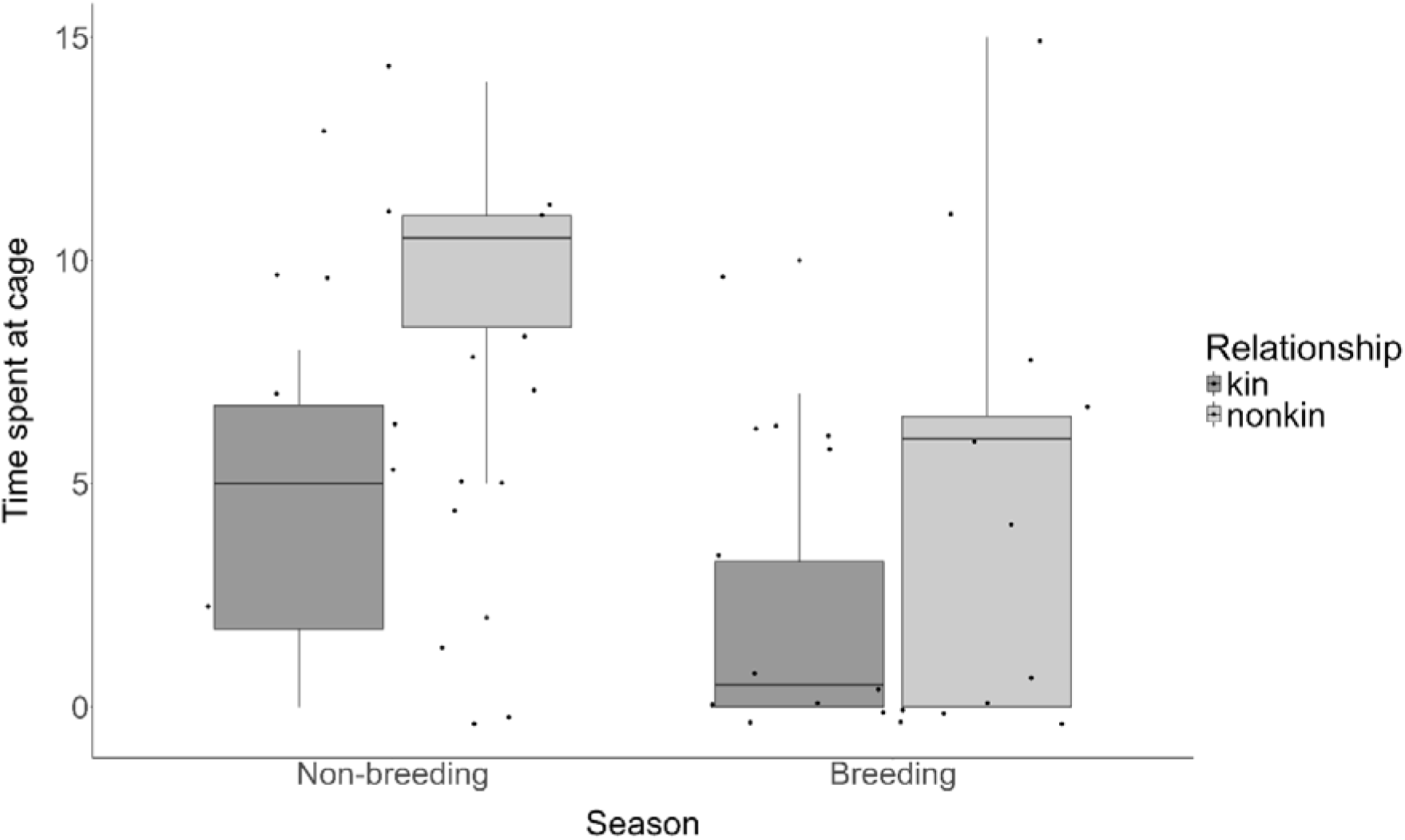
The time spent at the cage by focal females to neighbouring cage-housed bush Karoo rats during intruder tests. Boxplots show median and 1st and 3rd quartiles, the whiskers represent the minimum and maximum of the outlier data, and points represent individual values.

## Discussion

It is typically assumed that individuals of solitary species are intolerant of each other [42]. Here we tested whether female bush Karoo rats live solitarily because of high intra-specific aggression. However, we observed no aggression between mothers and their juvenile or even adult offspring. The mother-offspring relationship did not change when the offspring became older and had dispersed from their mother. In dyadic encounter tests, mother and offspring were regularly in body contact which did not decline as offspring became older. During field observations, mothers interacted rarely with pups and juveniles, but when they did so, it was mainly amicable. Thus, both experimental and field observation data indicate that aggression rarely occurs between mothers and offspring, aggression does not increase as offspring age and have become solitary, and is thus unlikely to be the mechanism leading to solitary living. This makes the alternative a likely explanation, i.e. motivation to disperse when reaching sexual maturity together with an absence of social attraction, such that dispersing individuals settle alone inside an unoccupied lodge.

It is typically assumed that aggression is the main mechanism leading to offspring dispersal and solitary living [43]. The association between aggression and dispersal in which subordinate individuals (juveniles) are driven out by dominant individuals (adults) is well known [43, 44]. However, there are equally many studies showing that aggression and dispersal are not always associated with one another [43]. In our study, mothers did not react more aggressively towards their dispersed offspring than towards their offspring still living with them. Female bush Karoo rats reach sexual maturity at six weeks [36] and are thus expected to leave their mother’s lodge (i.e. disperse) earliest at this age. Accordingly, females dispersed at 2 months of age but with a large variation that needs investigation in the future. For example, food availability, population density, and start vs end of breeding season are factors expected to influence dispersal. Our study showed the importance of dispersal in becoming solitary while there was no indication that aggression by the mother drives natal dispersal.

Aggressive interactions are often considered as the main underlying reason for solitary living and the reduction of aggression as a first step towards the evolution of sociality [45]. Dyadic encounters between adult neighbouring bush Karoo rats in a neutral arena were characterised by few interactions, independent of kinship, with sniffing as the predominant form of social investigation. Nearly no aggression occurred, not even between unrelated females. However, close kin were more likely to spend time in body contact with each other than non-kin. Thus, while female bush Karoo rats differentiated between kin and non-kin neighbours, neither relationship was driven by aggression. But there was also no social attraction between adults. We did however observe a high number of chases of striped mice by the bush Karoo rats, mostly in the breeding season (figure in supplementary file), demonstrating that bush Karoo rats can be aggressive. Within the same field site, and in similar experiments to ours, group-living striped mice were highly aggressive towards non-kin and interacted amicably with kin [46]. We had expected much more aggression in the solitary bush Karoo rat than in the sociable striped mice, and our ethical clearance protocol included that as soon as damaging fighting started, experiments would be terminated. We expected this to be the case regularly when non-kin met, but we never had to stop the encounter experiments. Therefore, absence of social attraction seems to be sufficient to lead to solitary living in bush Karoo rats, without the need for social intolerance and aggression.

Interactions between individuals are not random and instead reflect relatedness or familiarity [47]. In field intruder tests, where we expected to find more aggression towards conspecifics, due to the defence of resources [48], aggression was much more common during the breeding than the non-breeding season, and females were much more aggressive towards non-kin than kin. This indicates that female bush Karoo rats can differentiate between kin and non-kin, even when their kin had been living away from them (solitarily) for several months. Kin recognition thus occurs post-dispersal and indicates that mothers remember adult, dispersed offspring. This tells us that remembering kin (the mechanism for kin recognition) is persistent.

Female territoriality functions to defend resources and offspring [20, 49]. This theoretical consideration can explain why we observed nearly no aggression during dyadic encounter tests in a neutral test arena because no resources could be defended but did observe some during intruder tests when individuals defended their lodge. Does this indicate defence of resources such as food and shelter, or defence of offspring? More aggression was observed during the breeding season, which is also when more lodges are available due to low population density at the start of the season, and when food is highly abundant. Thus, our data support the female hypothesis of Wolff and Peterson [48] that territorial aggression is linked more to the defence of offspring (maternal aggression) than the defence of food resources. Reproductive competition can be high between female mammals [19], often leading to female infanticide [21], and is considered as one of the main reasons for solitary living [50]. Several other mammal species show maternal aggression, with females being aggressive at their nests in the breeding season while showing amicable behaviour at communal foraging grounds [reviewed by 48]. For example, female Arctic squirrels (*Urocitellus parryii* ) and grey squirrels (*Sciurus carolinensis* ) defend their territories near their nests but show considerable overlap in foraging areas. This behaviour is not limited to mammals but has been shown also in female social lizards (e.g., White’s skinks, *Egernia whitii*) where female aggression increased during pregnancy and after birth [49]. Aggression in female bush Karoo rats thus rather functions to protect their offspring than to establish a solitary social organisation.

## Conclusion

While solitary species have been considered to be generally asocial and aggressive, we have evidence that the solitary bush Karoo rat shows low levels of aggression. Instead, its social system is characterised by social tolerance, which suggests that lack of social attraction after dispersal and not aggression leads to solitary living. Female bush Karoo rats were not aggressive to their offspring even after they had dispersed from the maternal lodge and lived solitarily for months. In the solitary bush Karoo rat maternal aggression is not the mechanism driving solitary living. Tolerance of conspecifics is a condition for sociality. Therefore, solitary species that can tolerate their conspecifics may provide a model to understand the evolution of sociality in mammals.

## Ethical note

We adhered to the ASAB/ABS Guidelines for the Use of Animals in Research [51]. Bush Karoo rats were captured and handled using protocols approved by the Animal Ethics Screening Committee of the University of the Witwatersrand (AESC clearance number: 2018-03-15B). We further received additional ethical clearance for both the dyadic encounter tests and the field intruder presentation tests (AESC clearance numbers 2021/05/02/B and 22-12-024B respectively).

## Data availability

The data are available as supplementary material.

## Author contributions

L.M., C.S., and N.P. developed and designed the study. L.M., C.S. and S.P.S. collected the data. L.M. analysed the data, all authors discussed the results. L.M. and C.S. wrote the first draft of the manuscript. All co-authors contributed to the final draft of the manuscript.

## Acknowledgements

This study was made possible by the administrative and technical support of the Succulent Karoo Research Station (registered South African NPO 122-134). This study is part of the long-term Studies in Ecology and Evolution (SEE-Life) program of the CNRS.

---

**Figure S1.**
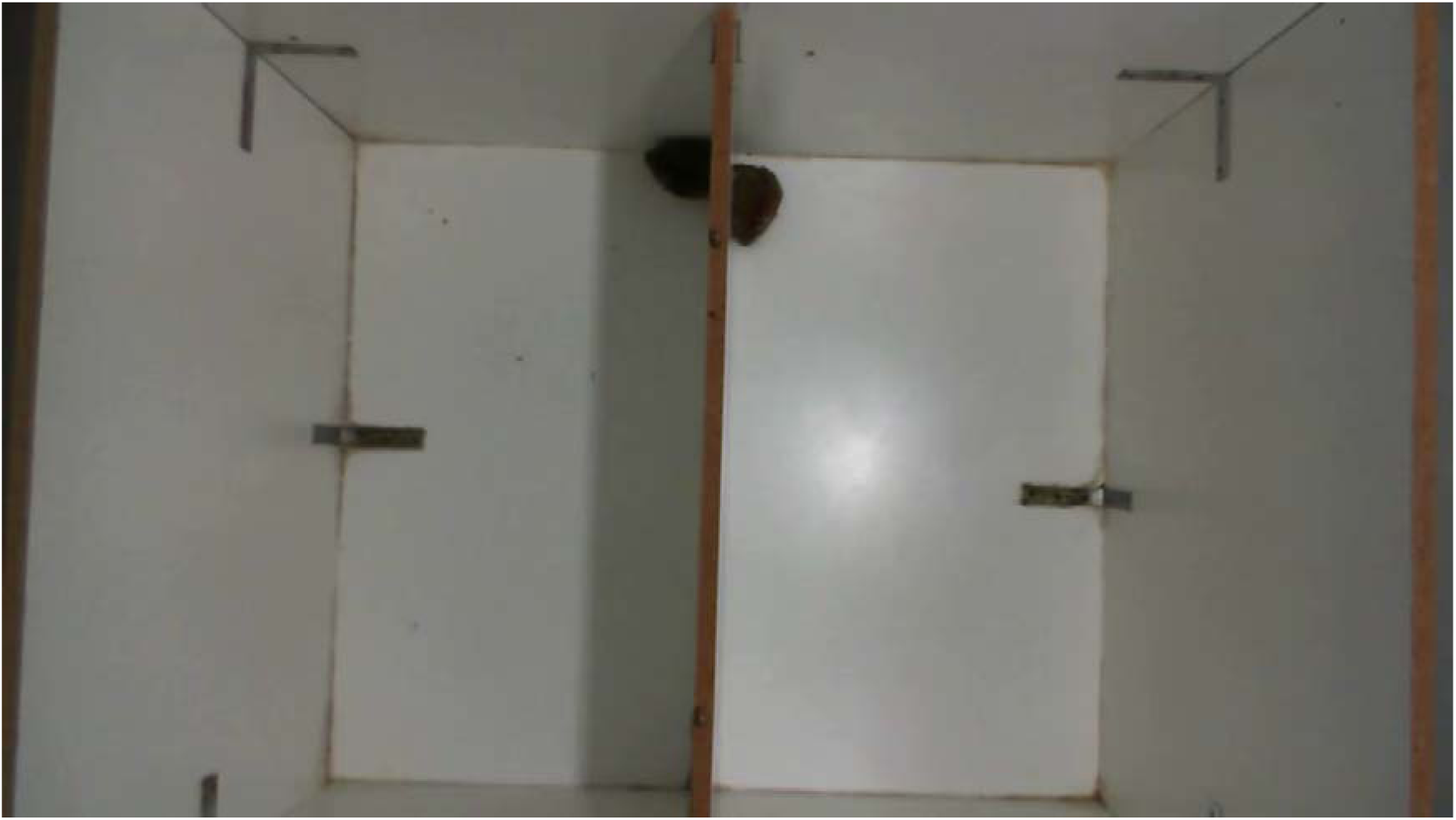
Picture showing the dyadic encounter test neutral arena with a partition in between. In the picture, two rats are seen sitting beside the partition.

**Figure S2.**
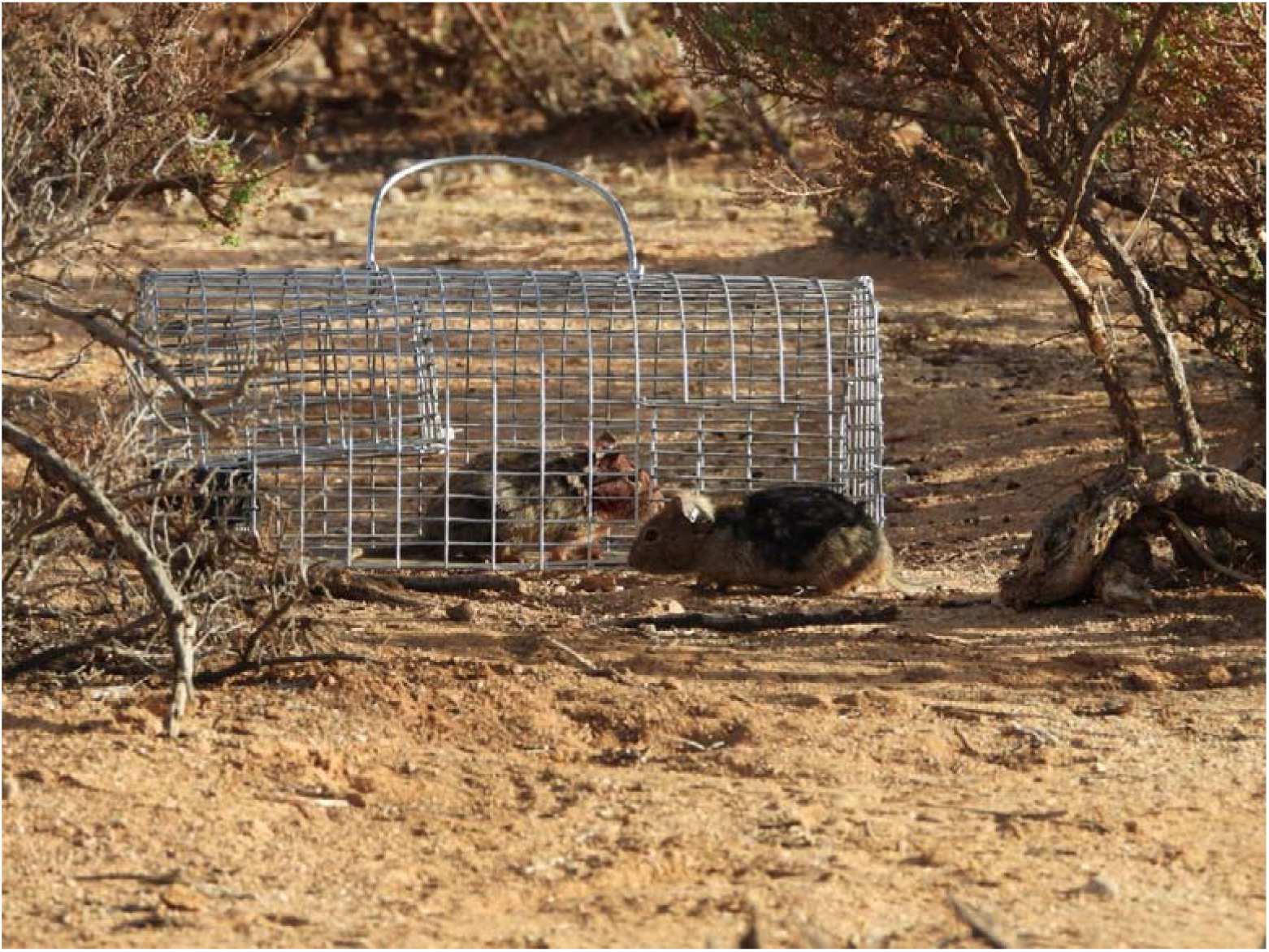
Figure showing the set-up for the field intruder tests. The focal rat is seen outside of the cage and the stimulus is inside the cage.

**Figure S3.**
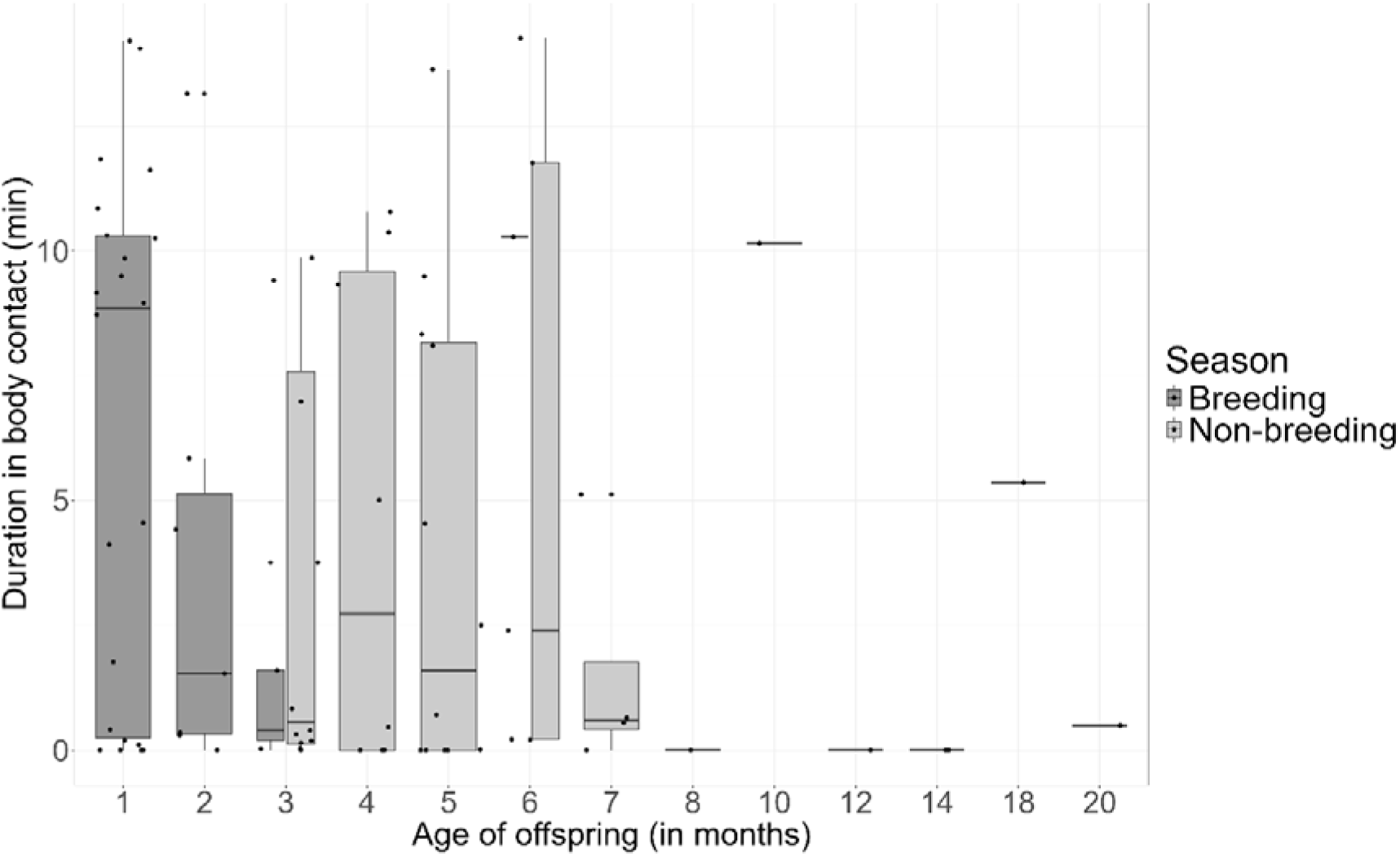
The duration spent in body contact during 15min observations by female bush Karoo rats according to the age of the offspring, for both seasons. Boxplots show median and 1st and 3rd quartiles, the whiskers represent the minimum and maximum of the outlier data and points represent individual values (breeding: n = 40, non-breeding: n = 39).

**Figure S4.**
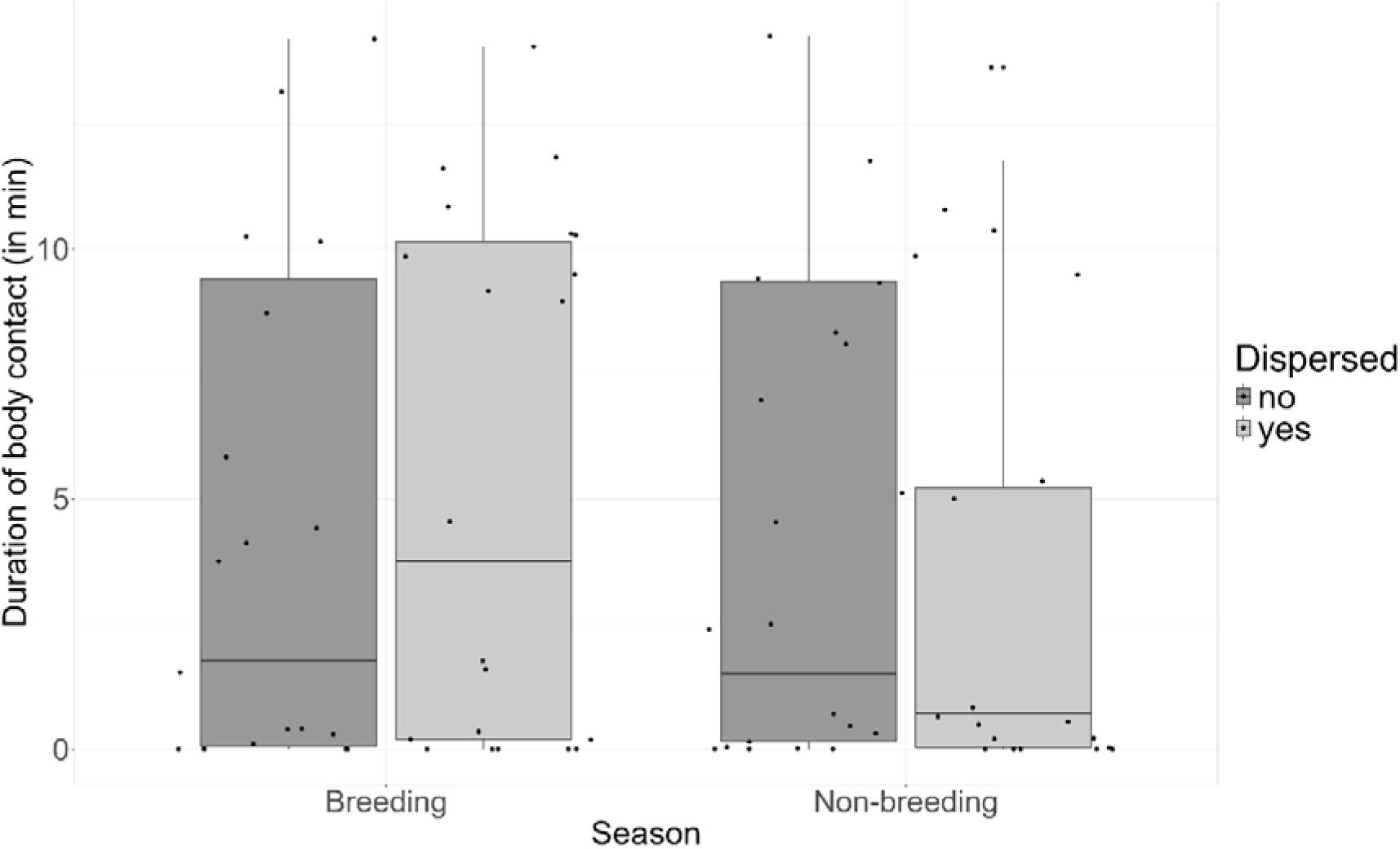
The duration spent in body contact by female bush Karoo rats according to whether offspring had dispersed or not, for both seasons. Boxplots show median and 1st and 3rd quartiles, the whiskers represent the minimum and maximum of the outlier data and points represent individual values (breeding: n = 40, non-breeding: n = 39).

**Figure S5.**
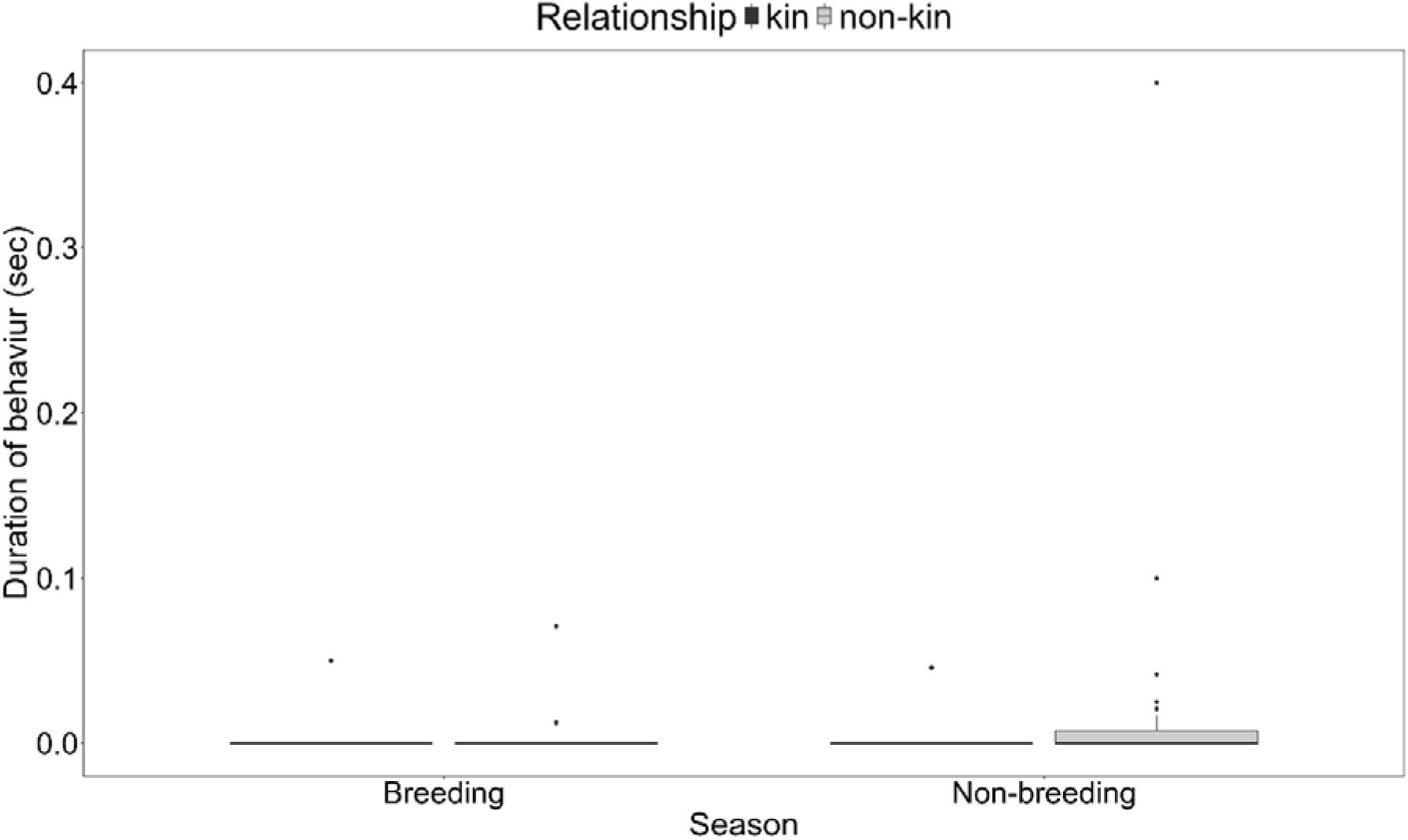
The duration spent fighting by female bush Karoo rats during dyadic encounter tests for both seasons. Boxplots show median and 1st and 3rd quartiles, the whiskers represent the minimum and maximum of the outlier data and points represent individual values (breeding: n = 40, non-breeding: n = 39).

**Figure S6.**
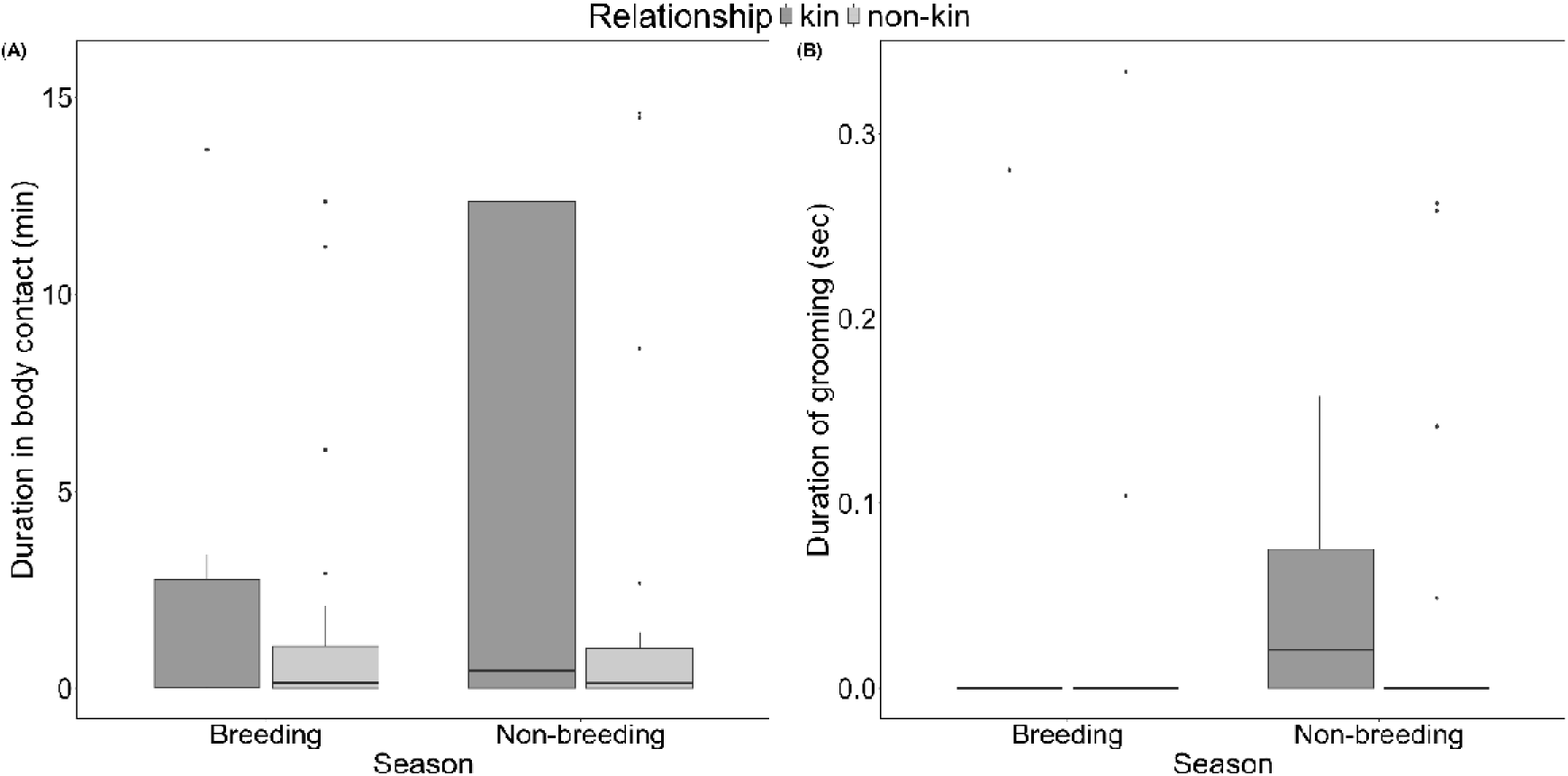
The duration of body contact (A) and grooming (B) behaviours between neighbouring female bush Karoo rats during dyadic encounter tests. Boxplots show median and 1st and 3rd quartiles, the whiskers represent the minimum and maximum of the outlier data and points represent individual values.

**Figure S7.**
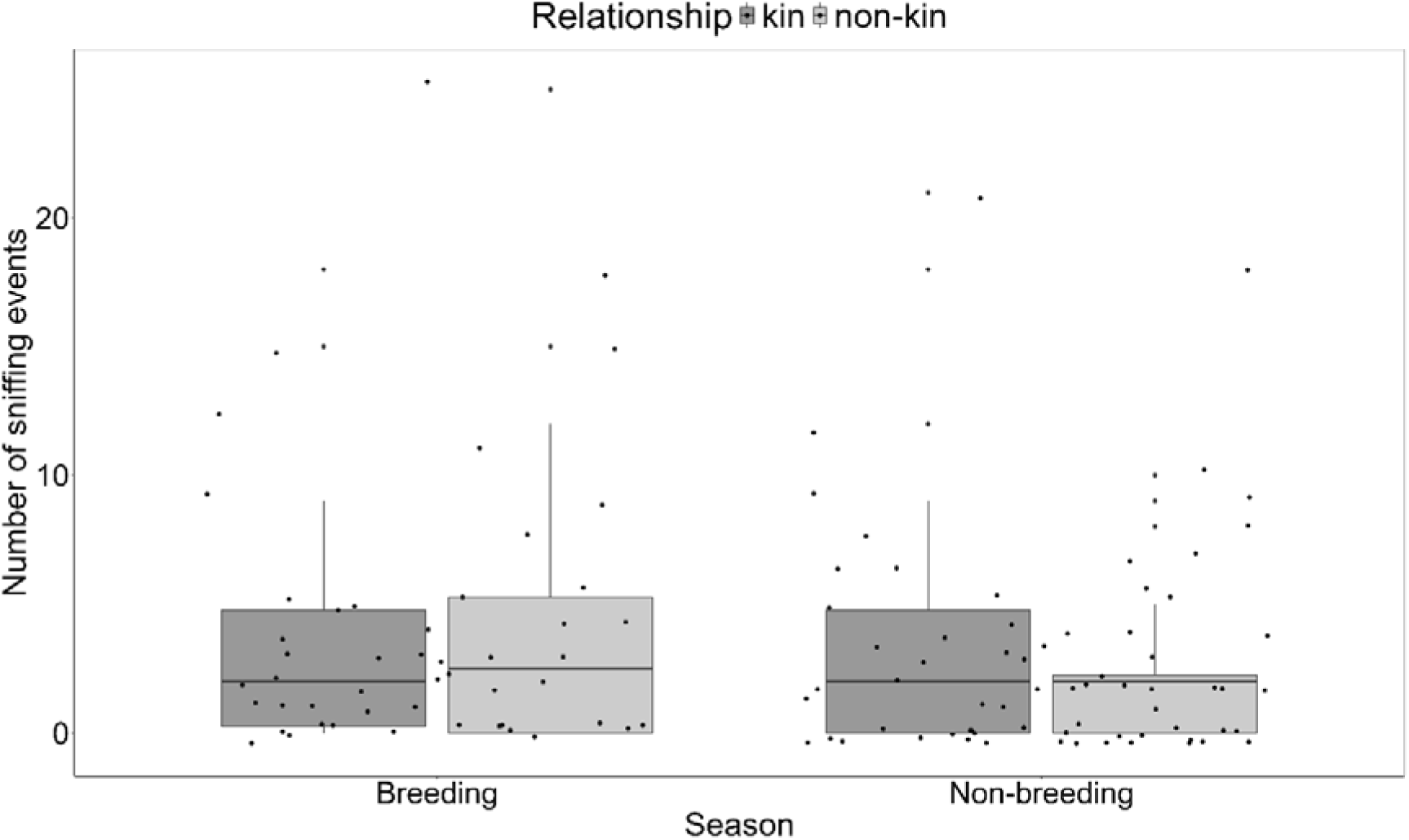
The number of sniffing events between neighbouring female bush Karoo rats during dyadic encounter tests. Boxplots show median and 1st and 3rd quartiles, the whiskers represent the minimum and maximum of the outlier data and points represent individual values.

**Figure S8.**
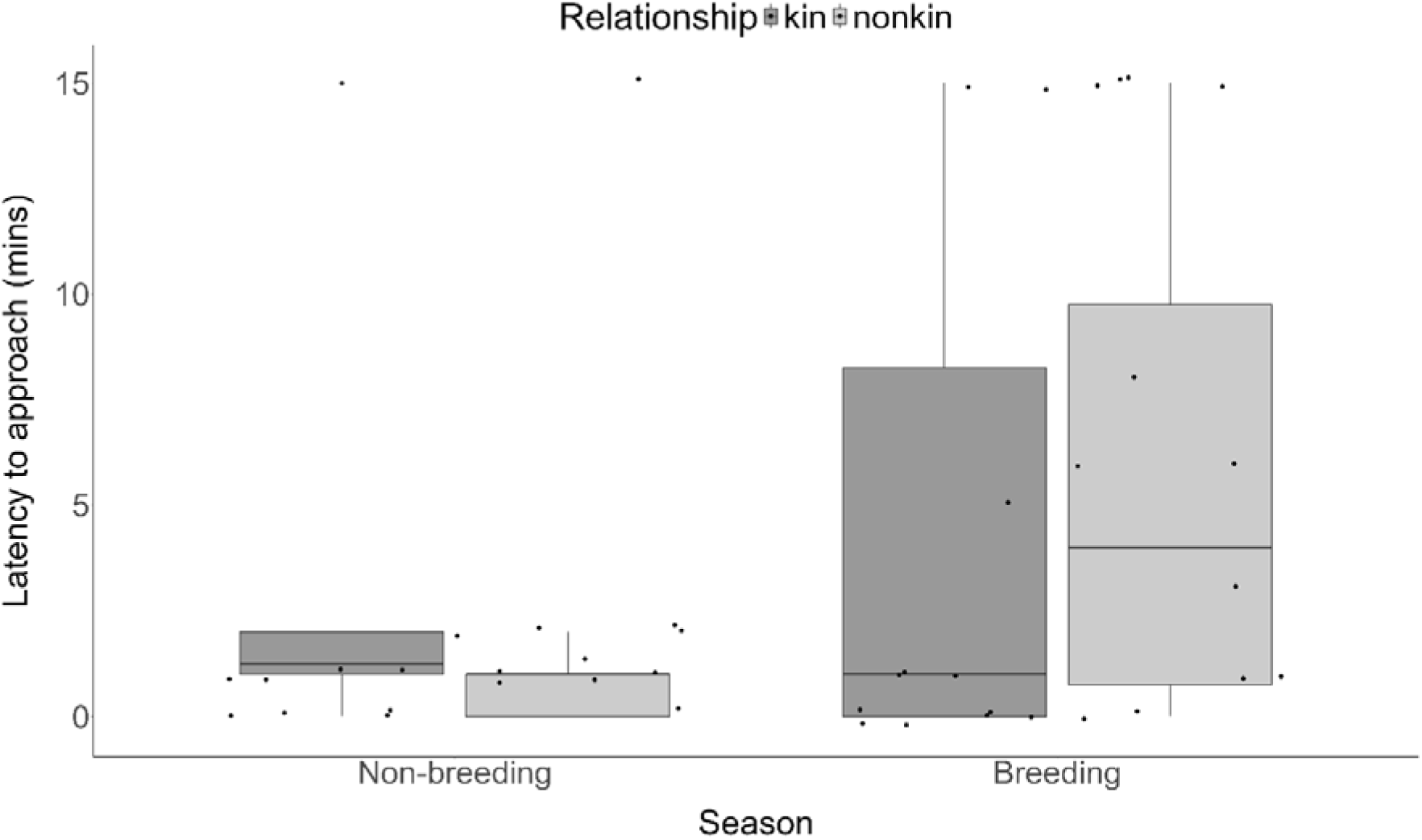
A measure of the latency to approach a cage during intruder tests based on the relationship between individuals. Boxplots show median and 1st and 3rd quartiles, the whiskers represent the minimum and maximum of the outlier data and points represent individual values.

**Figure S9.**
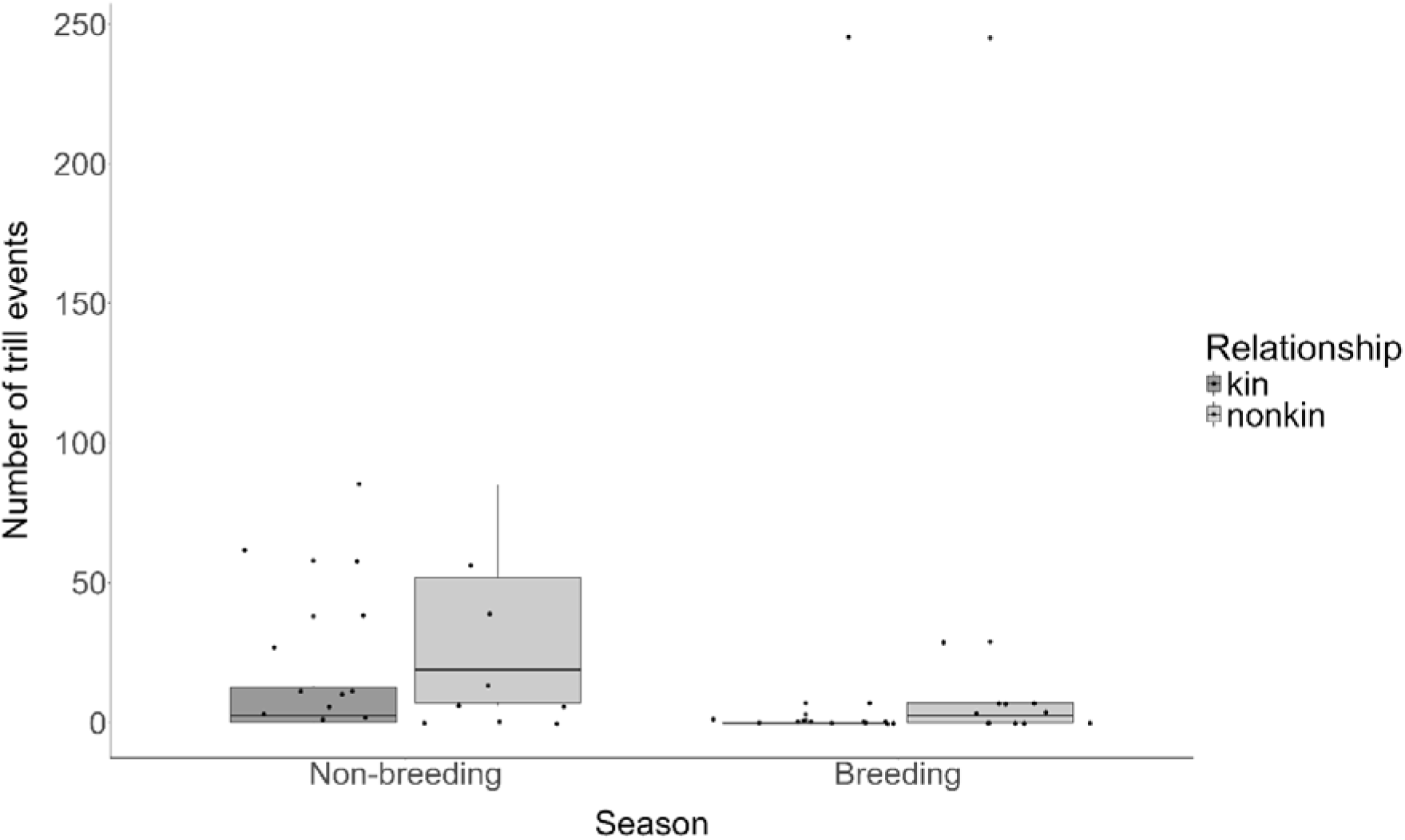
The number of trilling events measured during intruder tests. Boxplots show median and 1st and 3rd quartiles, the whiskers represent the minimum and maximum of the outlier data and points represent individual values.

**Figure S10.**
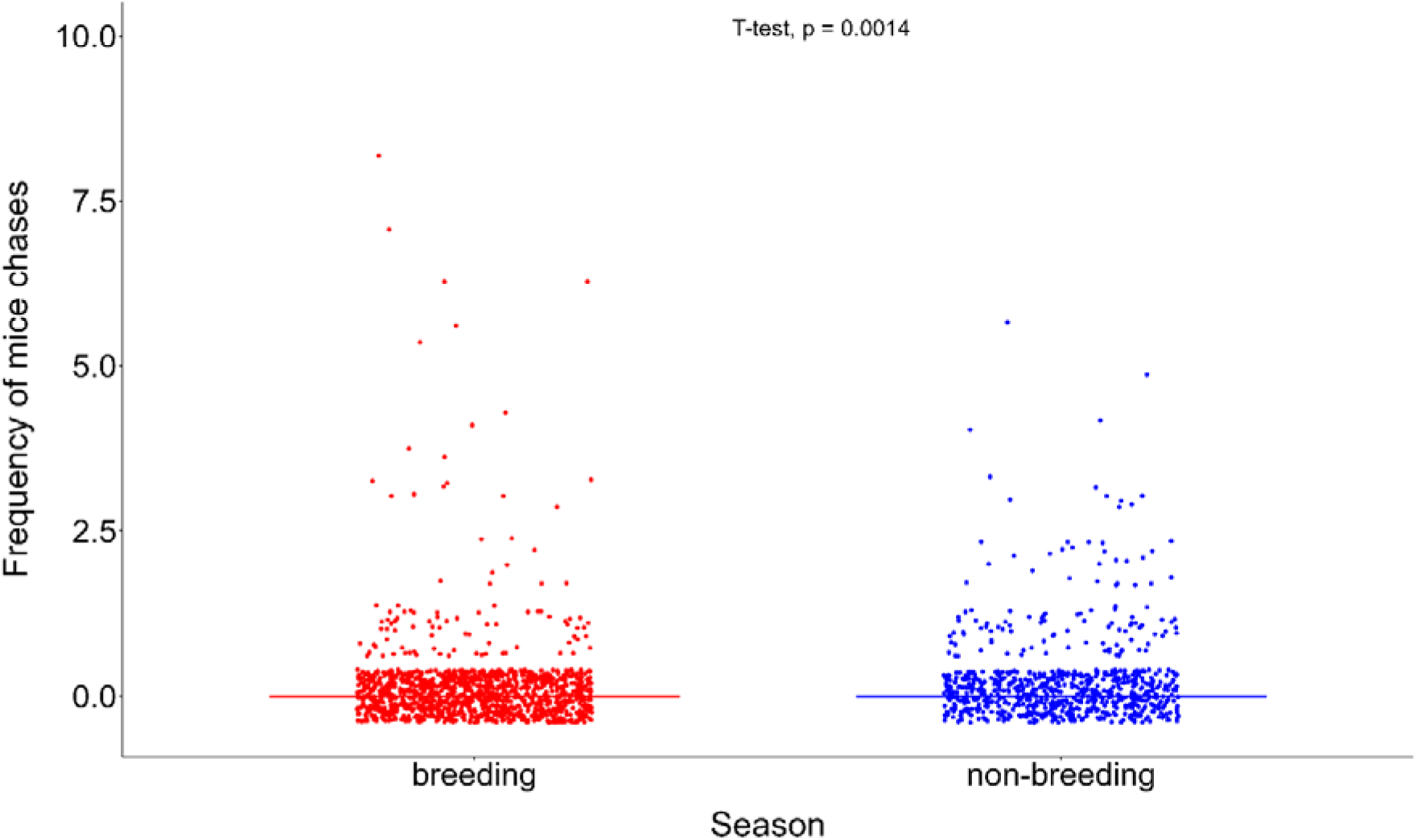
The frequency of chasing events against striped mice.

**Table S1:**
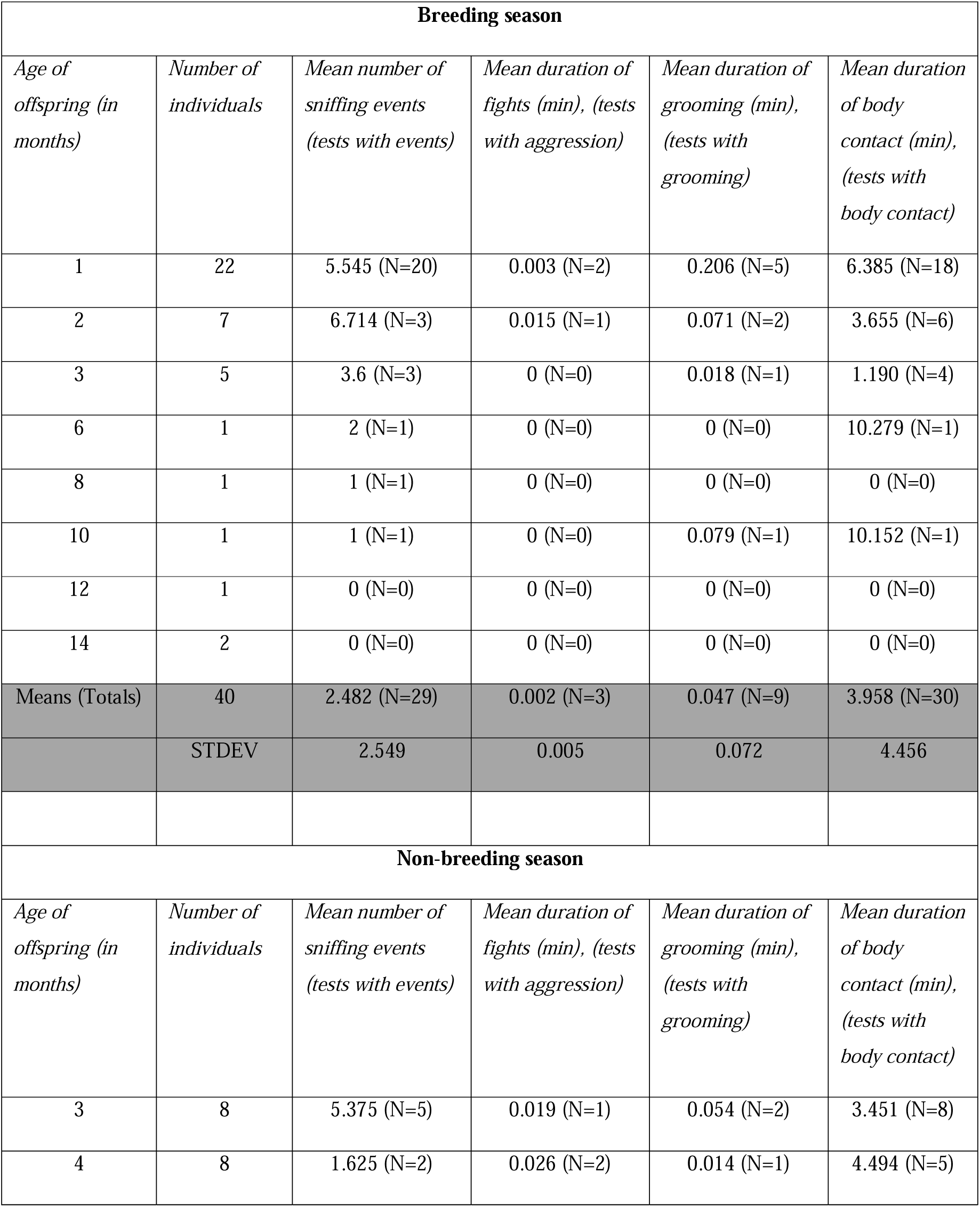
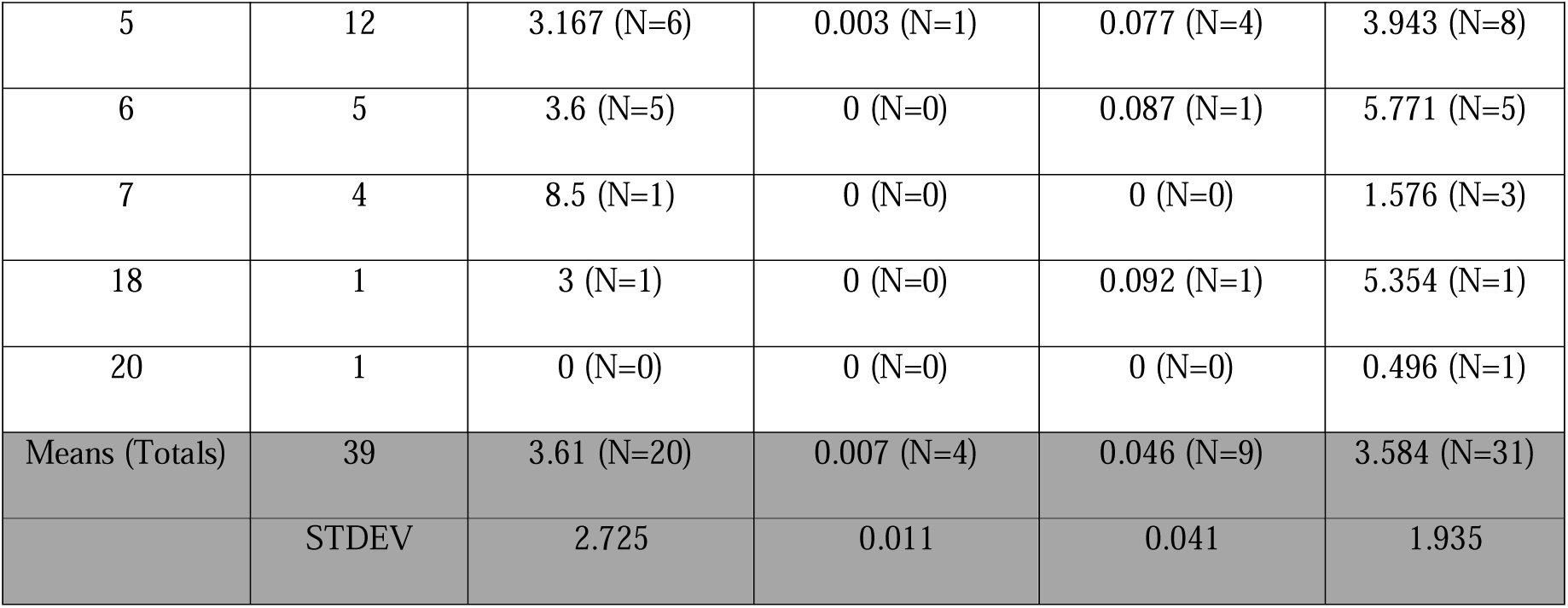
Sample sizes for the number of sniffing events and the time spent in fights, grooming and body contact between mothers and offspring during dyadic encounter tests grouped by season and the age of the offspring.

**Table S2:**
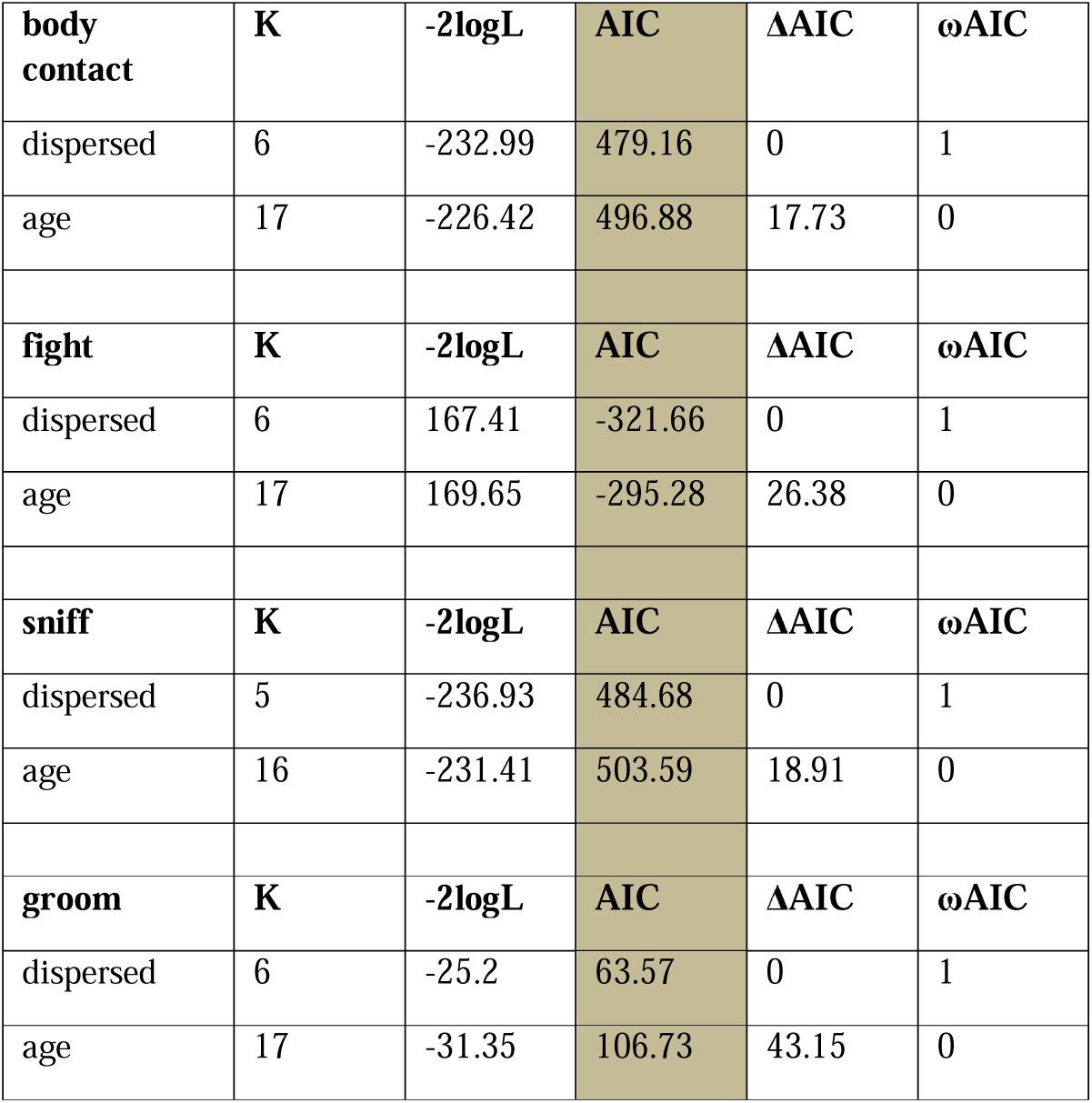
AIC-based model selection results examining reparameterization of age of offspring vs whether offspring had dispersed or not.

**Table S3.**
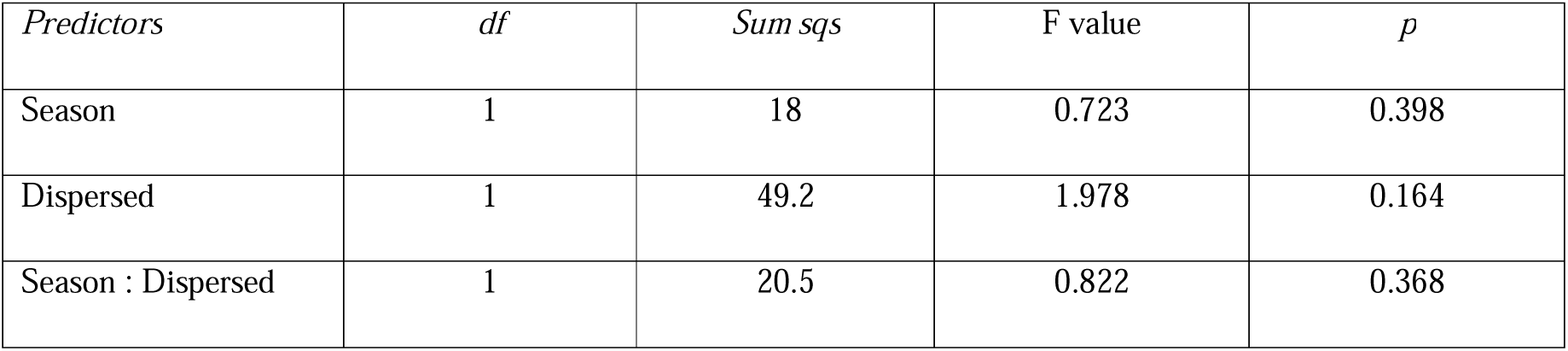
Sniffing: Results of the Anova models to test whether dispersal and the season have an impact on the frequency of sniffing events in the bush Karoo rat. *Sniff = season + dispersed*.

**Table S4.**
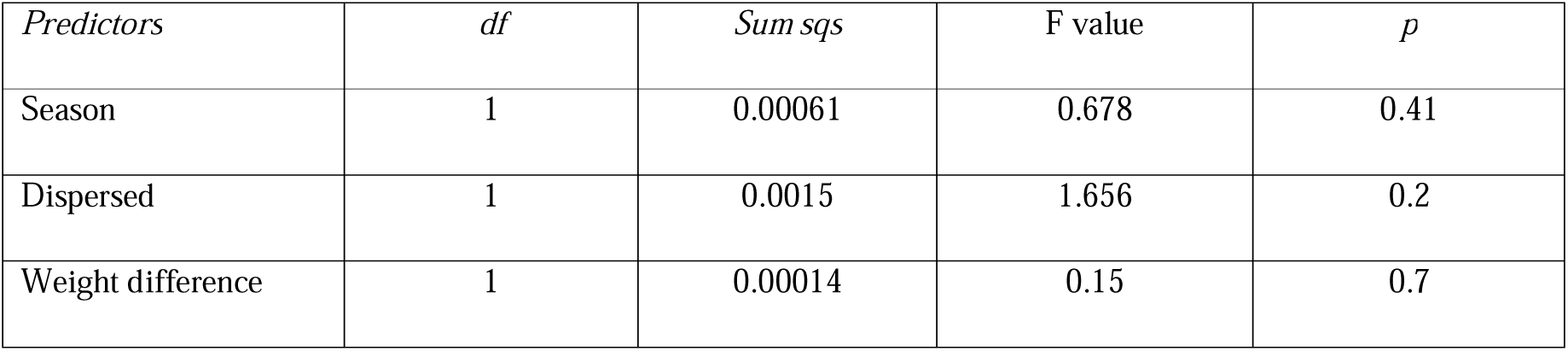
Fighting: Results of the Anova model to test whether dispersal and the season have an impact on the time spent fighting in the bush Karoo rat. *Fighting = season + dispersed*.

**Table S5.**
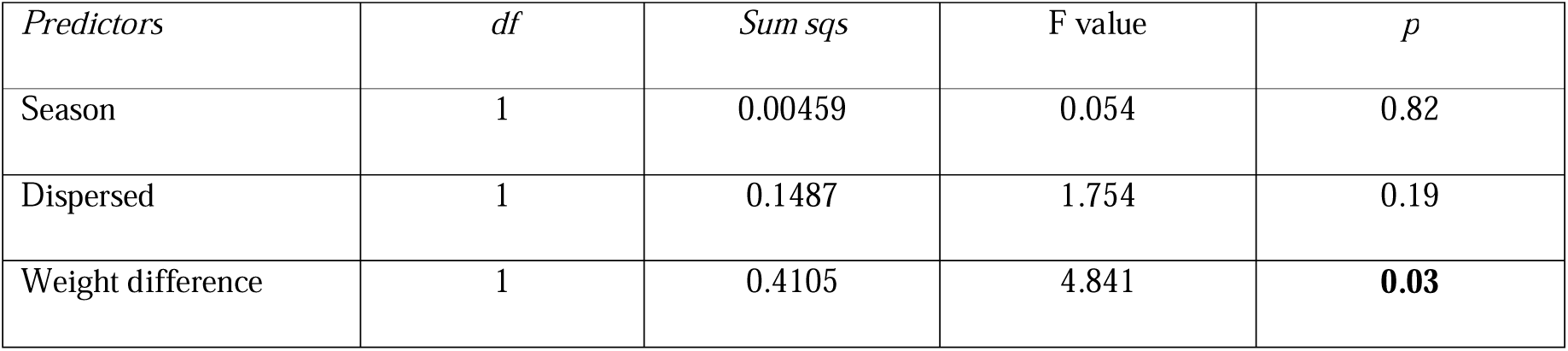
Grooming: Results of the Anova models to test whether dispersal and the season have an impact on the time spent grooming in the bush Karoo rat. Factors in bold type indicate significant predictors. *Grooming = season + dispersed*

**Table S6.**
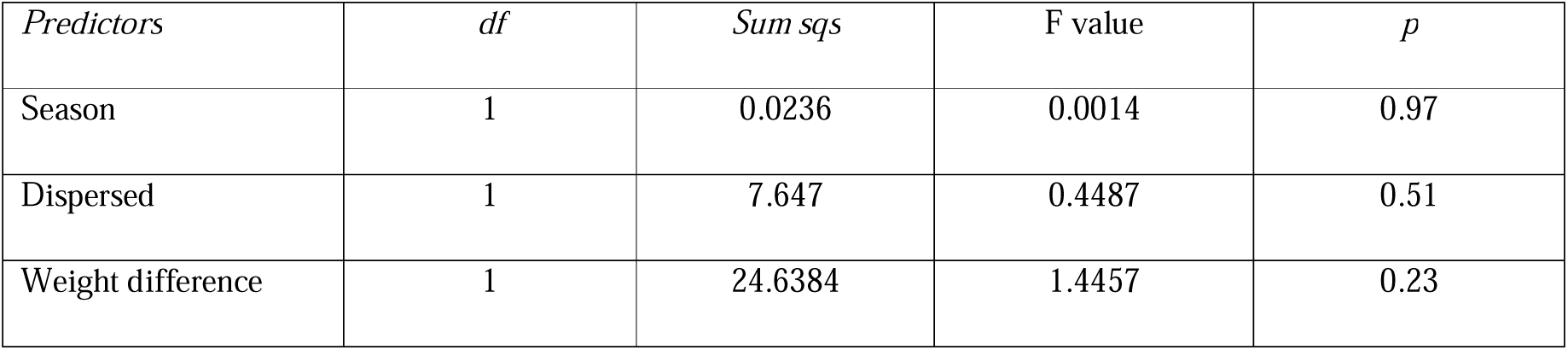
Body contact: Results of the Anova models to test whether dispersal and the season have an impact on the time spent in body contact in the bush Karoo rat. *Body contact = season + dispersed*

**Table S7.**
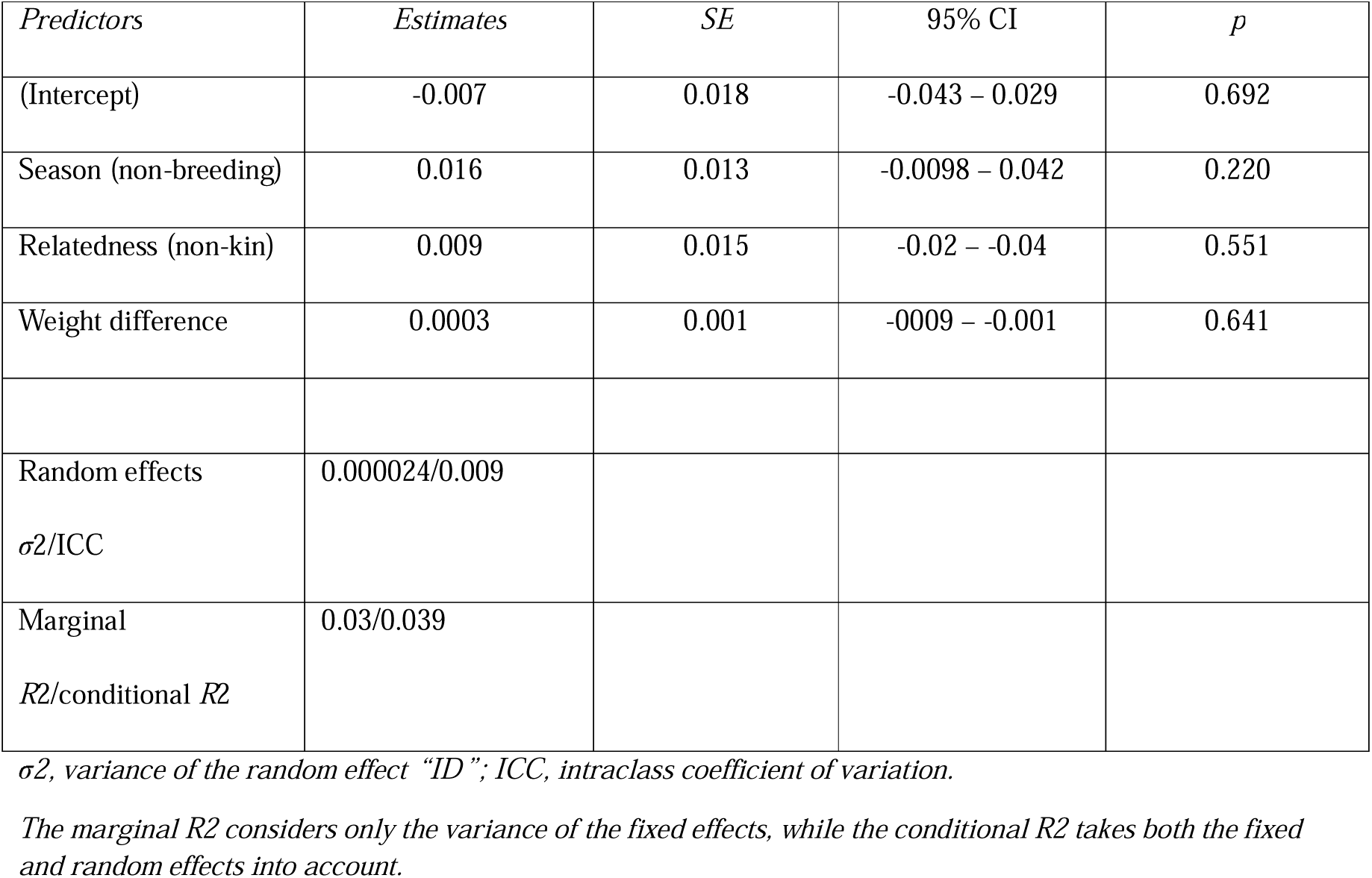
Neighbor encounter tests - Fighting: Results of the linear mixed effects models to identify which factors influenced time spent in amicable and aggressive behaviours by bush Karoo rats during dyadic encounter tests with neighbours. *Fight = season + relatedness + body size difference + ID-focal (random)*.

| *Predictors* | *Estimates* | *SE* | *95% CI* | *p* |
| --- | --- | --- | --- | --- |
| (Intercept) | -0.007 | 0.018 | -0.043 – 0.029 | 0.692 |
| Season (non-breeding) | 0.016 | 0.013 | -0.0098 – 0.042 | 0.220 |
| Relatedness (non-kin) | 0.009 | 0.015 | -0.02 – -0.04 | 0.551 |
| Weight difference | 0.0003 | 0.001 | -0.0009 – -0.001 | 0.641 |
| Random effects | 0.000024/0.009 |  |  |  |
| $\sigma^2$/ICC | | | | |
| Marginal | 0.03/0.039 |  |  |  |
| $R^2$/conditional $R^2$ | | | | |

$\sigma^2$ , variance of the random effect “ID”; ICC, intraclass coefficient of variation.
*The marginal $R^2$ considers only the variance of the fixed effects, while the conditional $R^2$ takes both the fixed* *and random effects into account.*

**Table S8.** Neighbor encounter tests – Body contact: Results of the linear mixed effects models to identify which factors influenced time spent in amicable and aggressive behaviours by bush Karoo rats during dyadic encounter tests with neighbours. Factors in bold type indicate significant predictors. *Body contact = season + relatedness + body size difference + ID-focal (random)*.

| *Predictors* | *Estimates* | *SE* | *95% CI* | *p* |
| --- | --- | --- | --- | --- |
| (Intercept) | 4.27 | 1.496 | 1.26 – 7.29 | < **0.01** |
| Season (non-breeding) | 0.57 | 1.094 | -1.63 – 2.80 | 0.60 |
| Relatedness (non-kin) | -2.36 | 1.316 | -5.04 – 0.31 | 0.08 |
| Weight difference | -0.02 | 0.048 | -0.12 – 0.08 | 0.69 |
| Random effects | 5.078/0.26 |  |  |  |
| $\sigma^2/ICC$ | | | | |
| Marginal | 0.06/0.305 |  |  |  |
| $R^2$/conditional $R^2$ | | | | |

$\sigma^2$ , variance of the random effect “ID”; ICC, intraclass coefficient of variation.
The marginal $R^2$ considers only the variance of the fixed effects, while the conditional $R^2$ takes both the fixed and random effects into account.

**Table S9.**
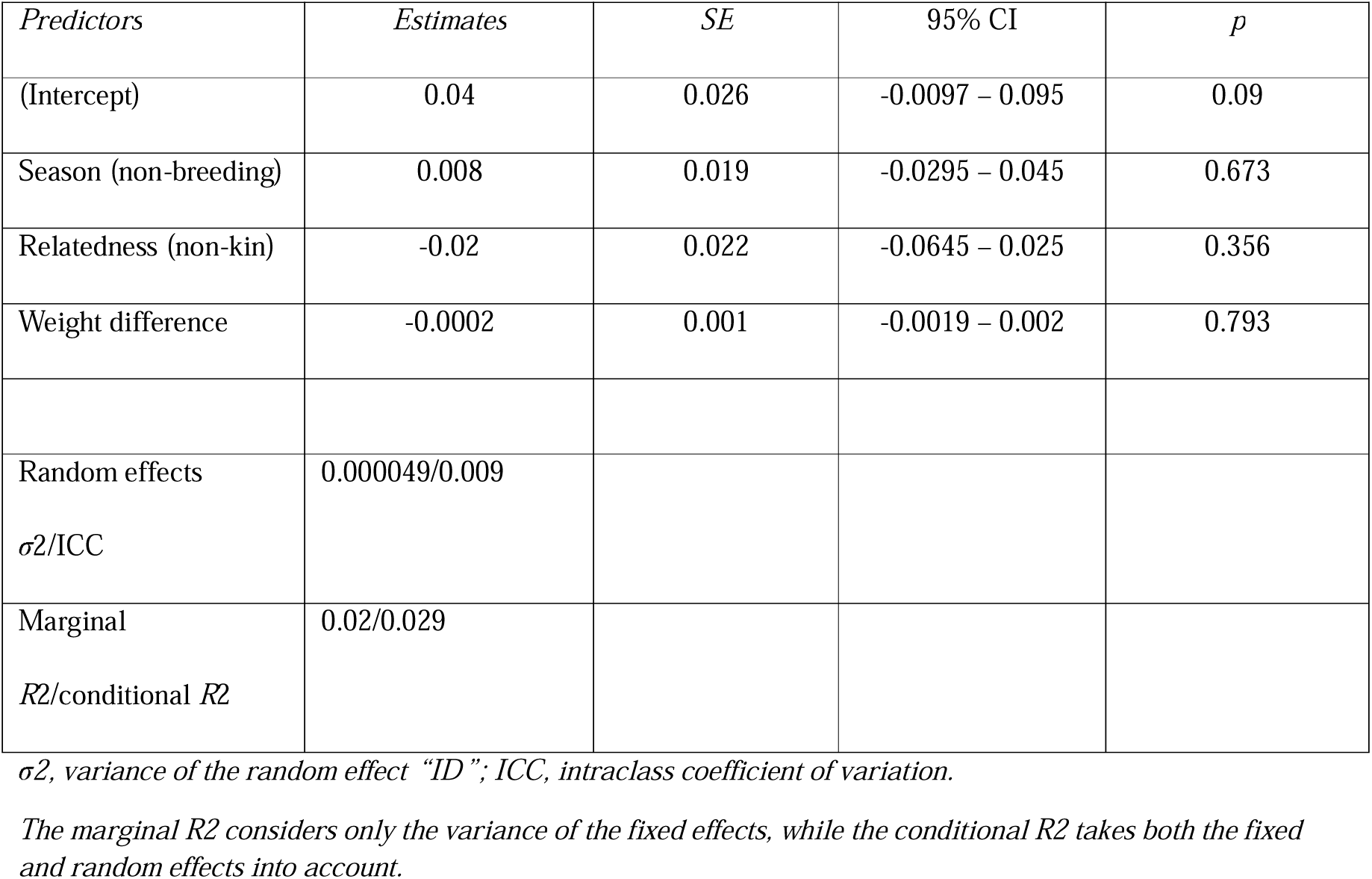
Neighbor encounter tests - Grooming: Results of the linear mixed effects models to identify which factors influenced time spent in amicable and aggressive behaviours by bush Karoo rats during dyadic encounter tests with neighbours. *Grooms = season + relatedness + body size difference + ID-focal (random)*.

| *Predictors* | *Estimates* | *SE* | *95% CI* | *p* |
| --- | --- | --- | --- | --- |
| (Intercept) | 0.04 | 0.026 | -0.0097 – 0.095 | 0.09 |
| Season (non-breeding) | 0.008 | 0.019 | -0.0295 – 0.045 | 0.673 |
| Relatedness (non-kin) | -0.02 | 0.022 | -0.0645 – 0.025 | 0.356 |
| Weight difference | -0.0002 | 0.001 | -0.0019 – 0.002 | 0.793 |
| Random effects | 0.000049/0.009 |  |  |  |
| $\sigma^2$/ICC | | | | |
| Marginal | 0.02/0.029 |  |  |  |
| $R^2$/conditional $R^2$ | | | | |

$\sigma^2$ , variance of the random effect “ID”; ICC, intraclass coefficient of variation.
The marginal $R^2$ considers only the variance of the fixed effects, while the conditional $R^2$ takes both the fixed and random effects into account.

**Table S10.**
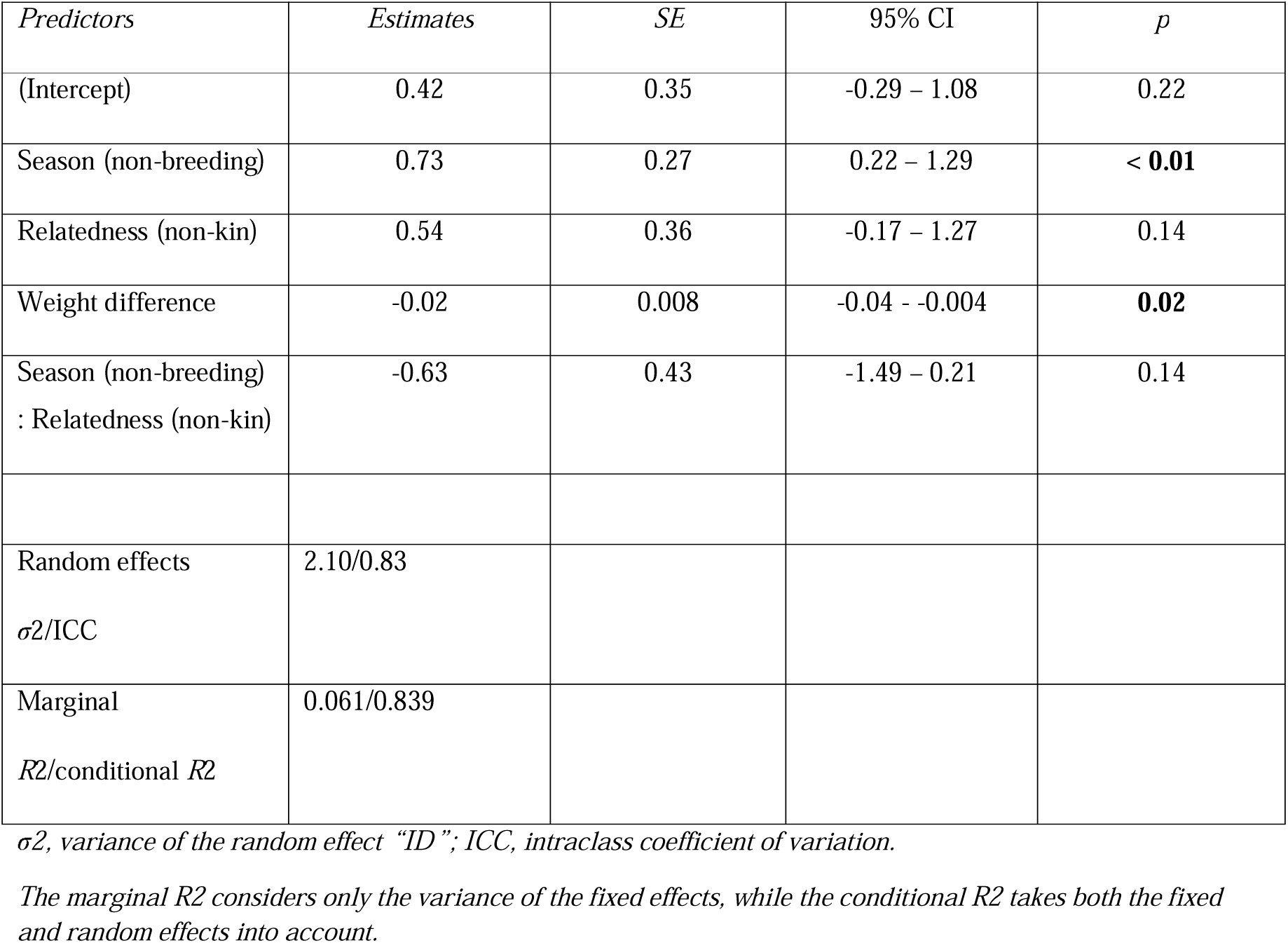
Sniffing events during encounter tests: Results of the generalized linear mixed effects models to identify which factors influenced the frequency of sniffing events exhibited by bush Karoo rats during dyadic encounter tests towards neighbours. Factors in bold type indicate significant predictors. *Sniff = season * relatedness + body size difference + ID-focal (random) + ID-stimulus(random)*.

**Table S11.**
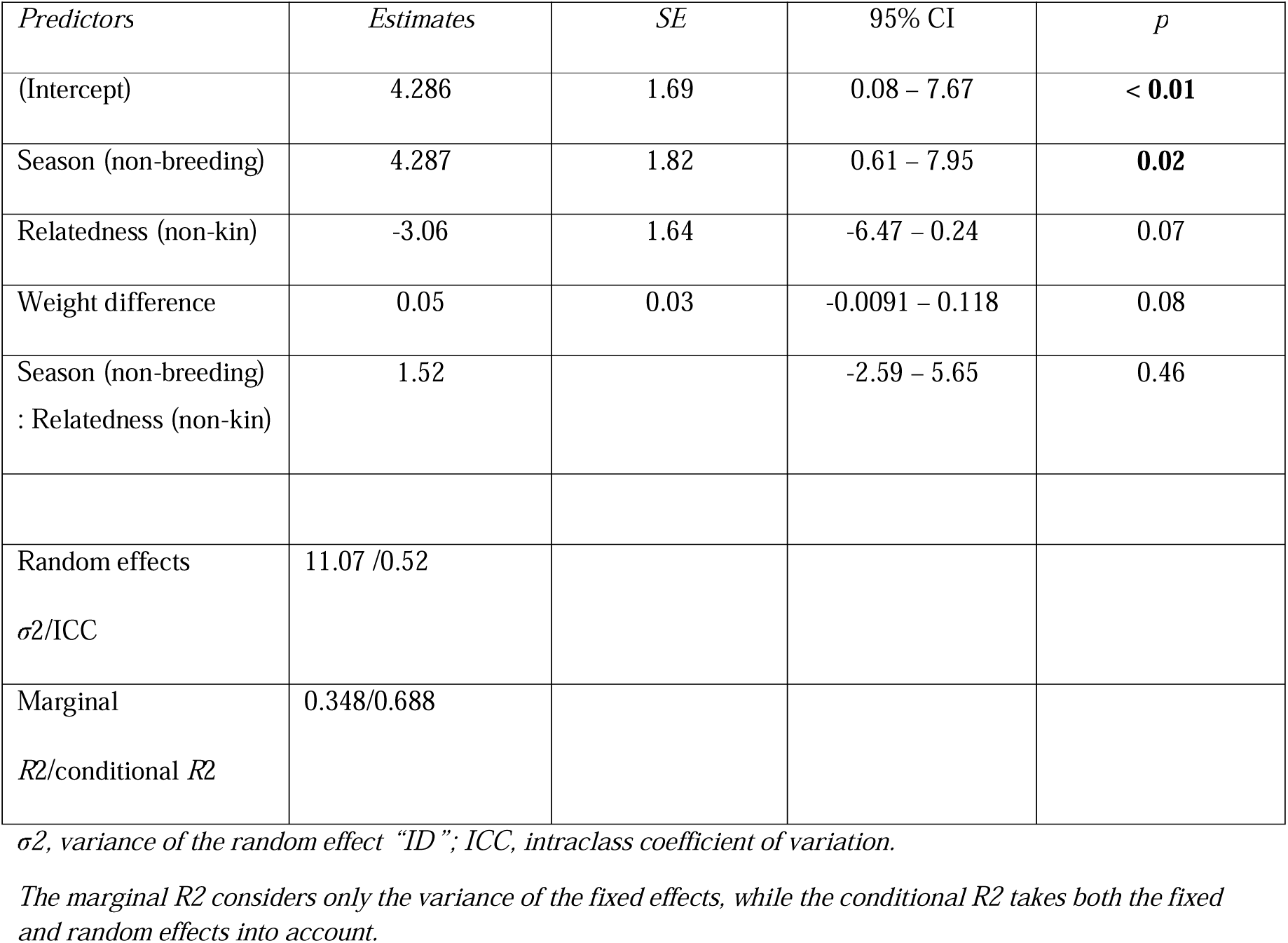
Latency to first aggression during intruder tests: Results of the linear mixed effects models to identify which factors influenced the latency to the first aggressive behaviour with a stimulus during intruder tests towards neighbours. *Latency to first aggression = season * relatedness + body size difference + ID-focal (random)*.

**Table S12.**
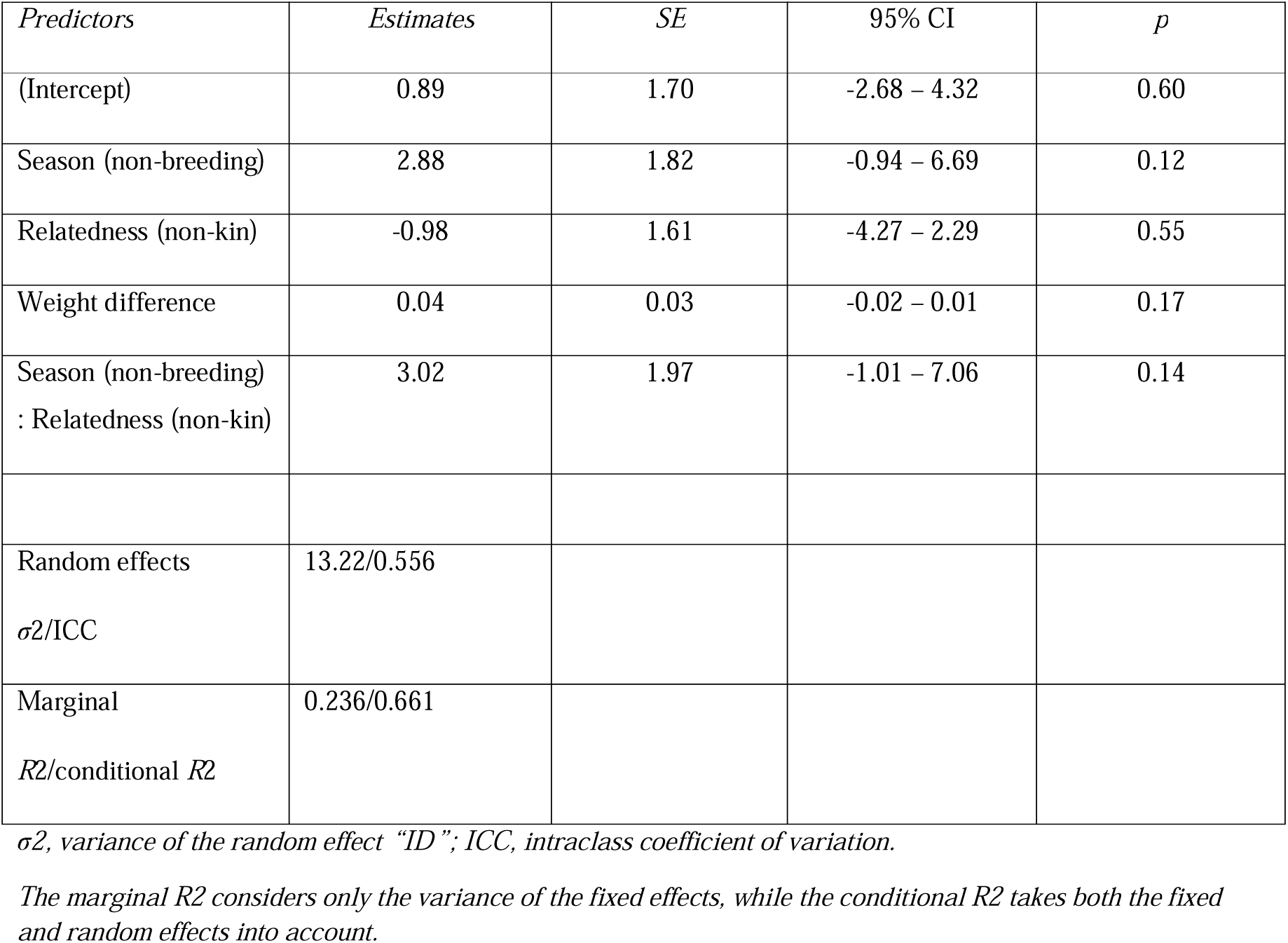
Latency to approach cage during intruder tests: Results of the linear mixed effects models to identify which factors influenced the latency to approach a cage with a stimulus during intruder tests towards neighbours. *Latency to approach = season * relatedness + body size difference + ID-focal (random)*.

**Table S13.**
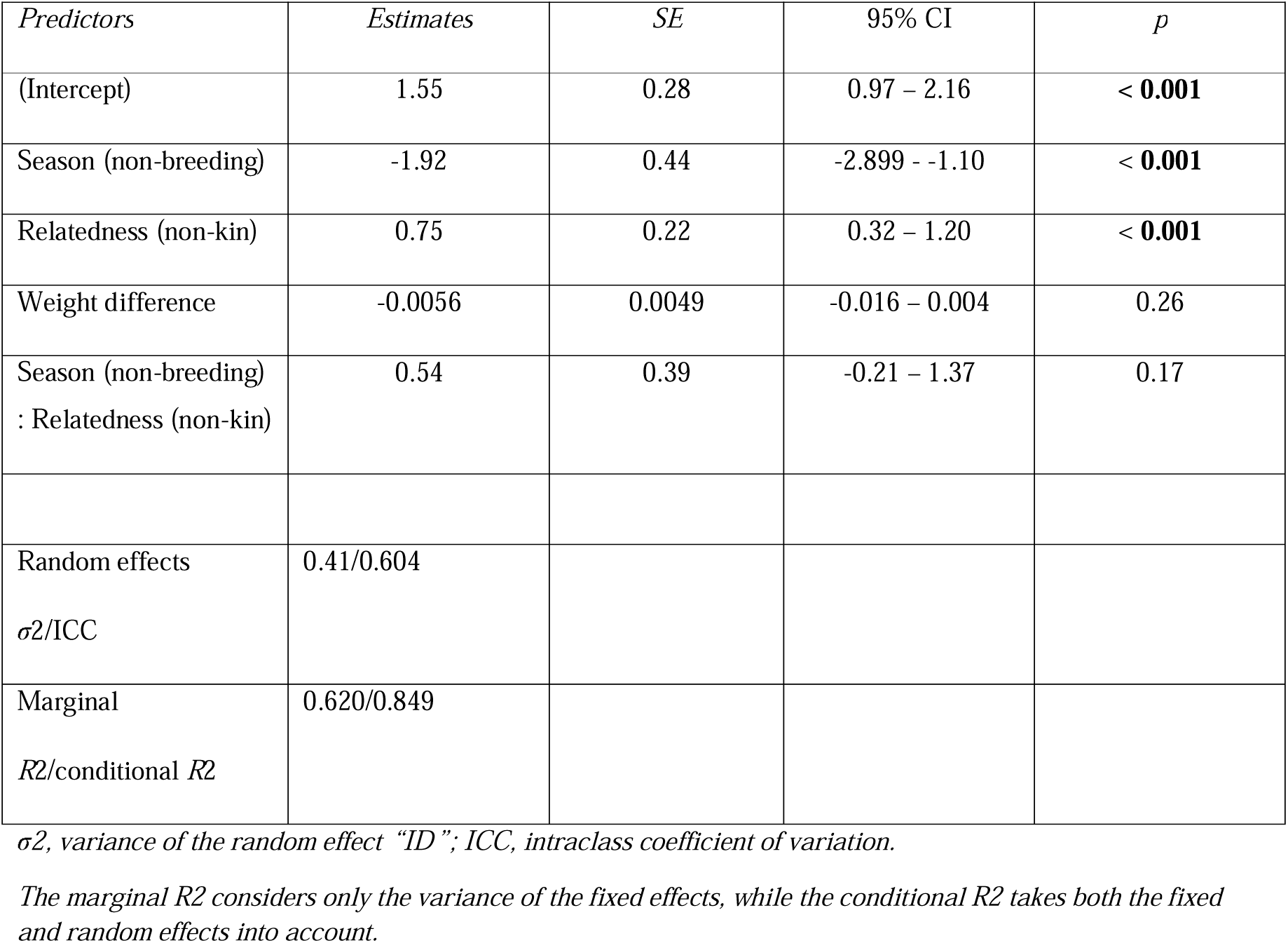
Charging events during intruder tests: Results of the generalized linear mixed effects models to identify which factors influenced the frequency of charging events exhibited by bush Karoo rats during intruder tests towards neighbours. Factors in bold type indicate significant predictors. *Charging = season * relatedness + body size difference + ID-focal (random)*.

**Table S14.**
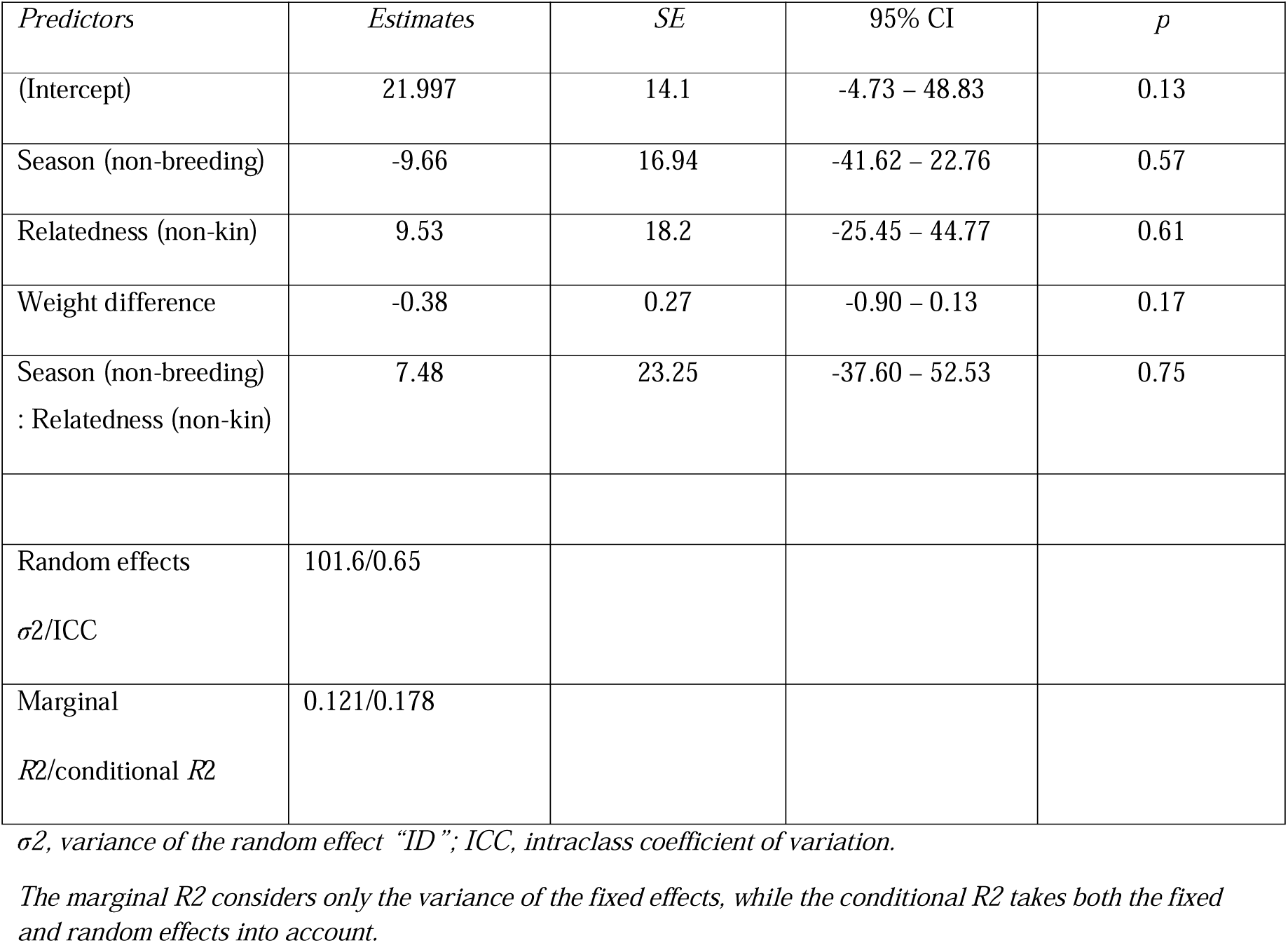
Number of trill sounds during intruder tests: Results of the linear mixed effects models to identify which factors influenced the number of trill sounds with a stimulus during intruder tests towards neighbours. *Trills = season * relatedness + body size difference + ID-focal (random)*.

| *Predictors* | *Estimates* | *SE* | *95% CI* | *p* |
| --- | --- | --- | --- | --- |
| (Intercept) | 21.997 | 14.1 | -4.73 – 48.83 | 0.13 |
| Season (non-breeding) | -9.66 | 16.94 | -41.62 – 22.76 | 0.57 |
| Relatedness (non-kin) | 9.53 | 18.2 | -25.45 – 44.77 | 0.61 |
| Weight difference | -0.38 | 0.27 | -0.90 – 0.13 | 0.17 |
| Season (non-breeding)<br>: Relatedness (non-kin) | 7.48 | 23.25 | -37.60 – 52.53 | 0.75 |
| Random effects | 101.6/0.65 |  |  |  |
| $\sigma^2/\text{ICC}$ | | | | |
| Marginal<br>$R^2/\text{conditional } R^2$ | 0.121/0.178 | | | |

$\sigma^2$ , variance of the random effect “ID”; ICC, intraclass coefficient of variation.
The marginal $R^2$ considers only the variance of the fixed effects, while the conditional $R^2$ takes both the fixed and random effects into account.

**Table S15.**
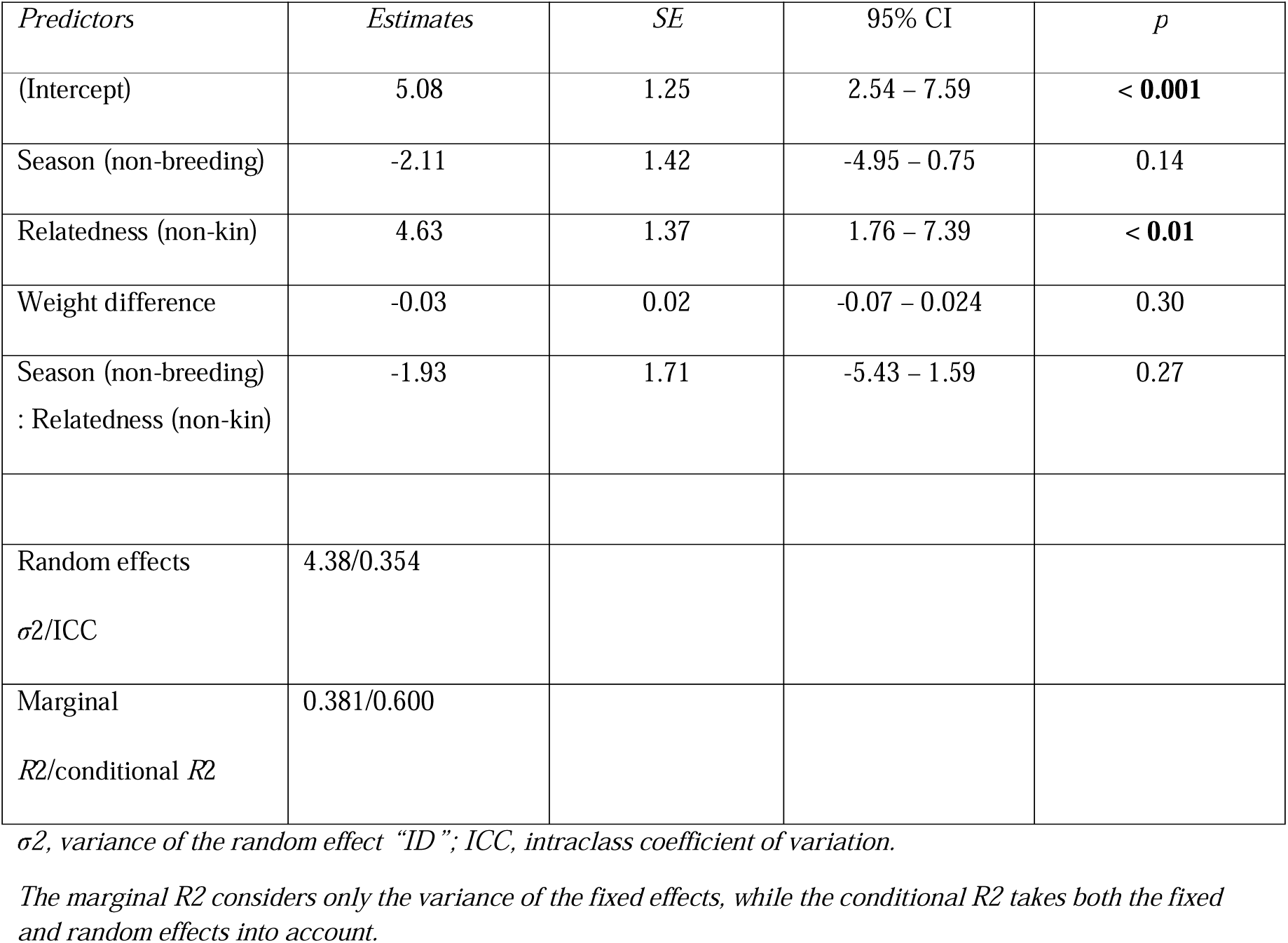
Time spent at a cage during intruder tests: Results of the linear mixed effects models to identify which factors influenced the time spent at a cage of the stimulus during intruder tests. Factors in bold type indicate significant predictors. *Time at cage = season * relatedness + body size difference + ID-focal (random)*.

**Table S3.1.**
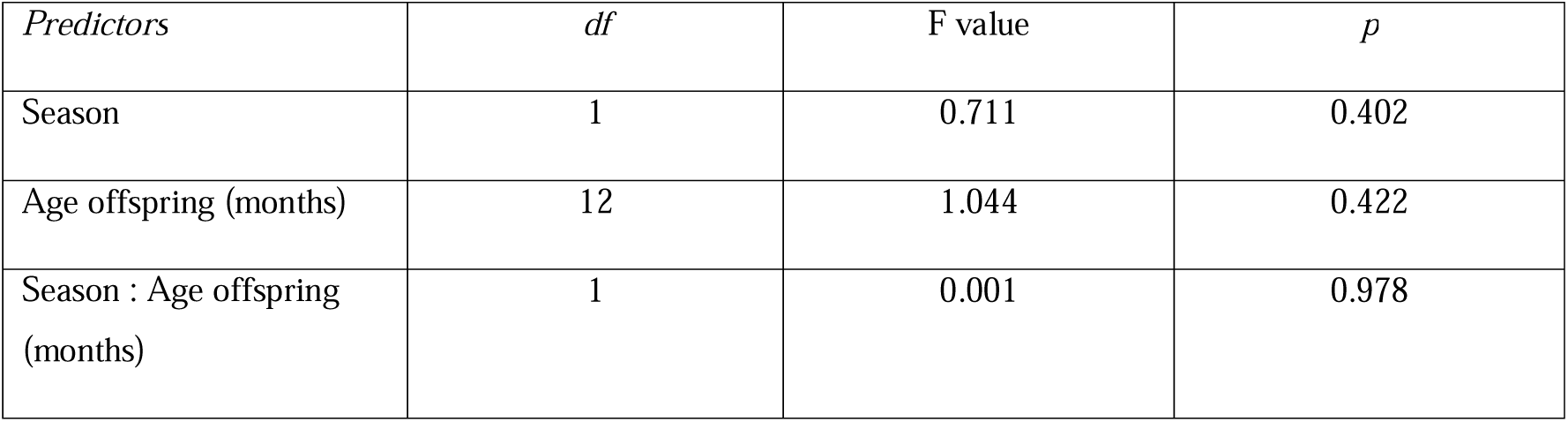
1. Sniffing: Results of the Anova models to test whether age of offspring and the season have an impact on the frequency of sniffing events in the bush Karoo rat. *Sniff = season + age of offspring*

**Table S4.1.**
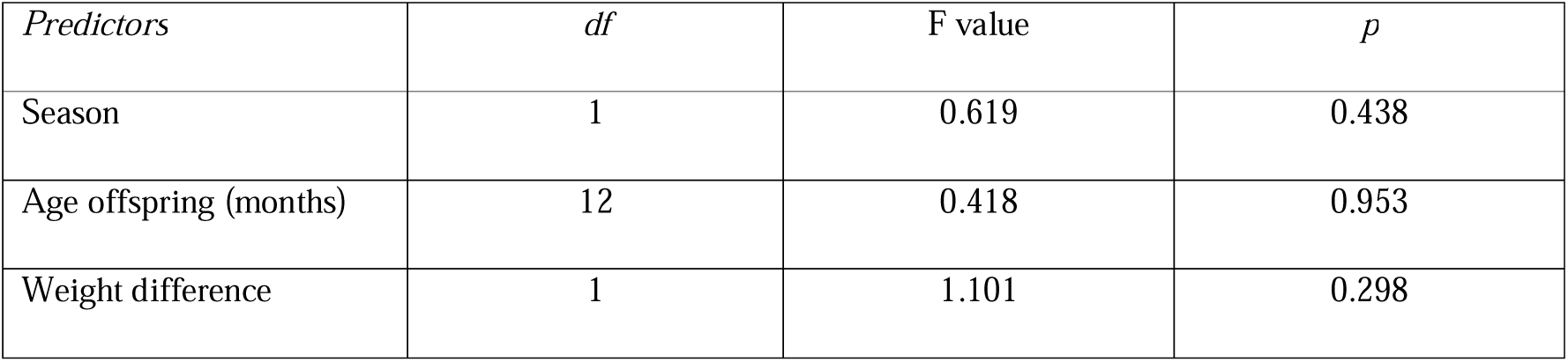
Fighting: Results of the Anova model to test whether age of offspring and the season have an impact on the time spent fighting in the bush Karoo rat. *Fighting = season + age of offspring*

**Table S5.1.**
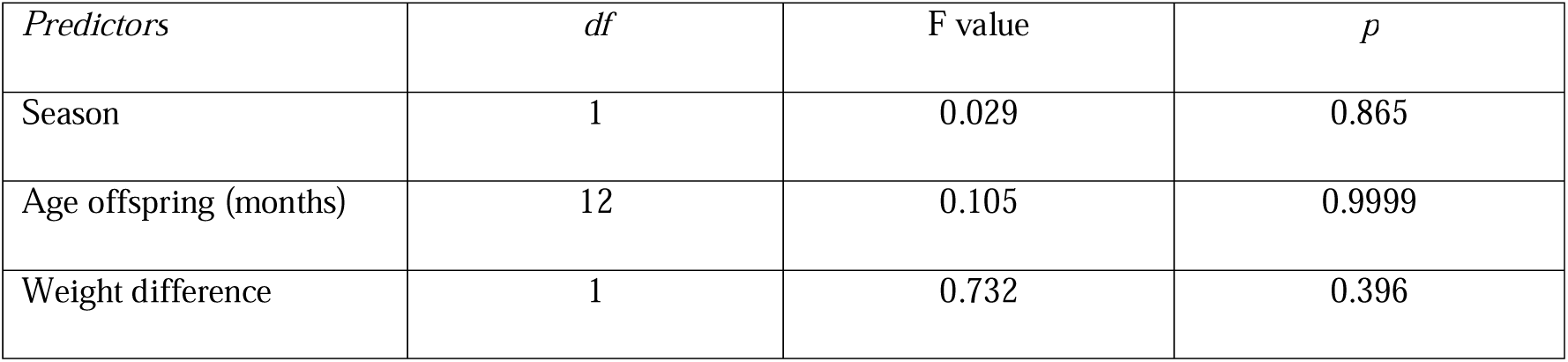
Grooming: Results of the Anova models to test whether age of offspring and the season have an impacton the time spent grooming in the bush Karoo rat. *Grooming = season + age of offspring*

**Table S6.1.**
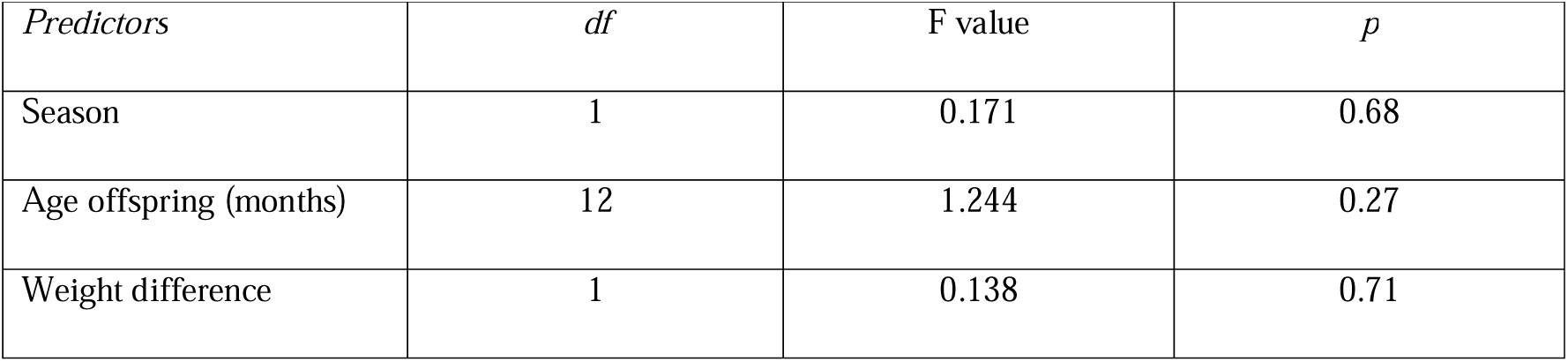
Body contact: Results of the Anova models to test whether age of offspring and the season have an impact on the time spent in body contact in the bush Karoo rat. *Body contact = season + age of offspring*

