## Supplementary_file for "Tolerant mothers: aggression does not explain solitary living in the bush Karoo rat"

**Electronic supplementary file**

**Supplementary tables**

**Table S1**: Sample sizes for the number of sniffing events and the time spent in fights, grooming and body contact between mothers and offspring during dyadic encounter tests grouped by season and the age of the offspring.

| **Breeding season** | | | | | |
| --- | --- | --- | --- | --- | --- |
| *Age of offspring (in months)* | *Number of individuals* | *Mean number of sniffing events (tests with events)* | *Mean duration of fights (min), (tests with aggression)* | *Mean duration of grooming (min), (tests with grooming)* | *Mean duration of body contact (min), (tests with body contact)* |
| 1 | 22 | 5.545 (N=20) | 0.003 (N=2) | 0.206 (N=5) | 6.385 (N=18) |
| 2 | 7 | 6.714 (N=3) | 0.015 (N=1) | 0.071 (N=2) | 3.655 (N=6) |
| 3 | 5 | 3.6 (N=3) | 0 (N=0) | 0.018 (N=1) | 1.190 (N=4) |
| 6 | 1 | 2 (N=1) | 0 (N=0) | 0 (N=0) | 10.279 (N=1) |
| 8 | 1 | 1 (N=1) | 0 (N=0) | 0 (N=0) | 0 (N=0) |
| 10 | 1 | 1 (N=1) | 0 (N=0) | 0.079 (N=1) | 10.152 (N=1) |
| 12 | 1 | 0 (N=0) | 0 (N=0) | 0 (N=0) | 0 (N=0) |
| 14 | 2 | 0 (N=0) | 0 (N=0) | 0 (N=0) | 0 (N=0) |
| Means (Totals) | 40 | 2.482 (N=29) | 0.002 (N=3) | 0.047 (N=9) | 3.958 (N=30) |
|  | STDEV | 2.549 | 0.005 | 0.072 | 4.456 |
| **Non-breeding season** | | | | | |
| *Age of offspring (in months)* | *Number of individuals* | *Mean number of sniffing events (tests with events)* | *Mean duration of fights (min), (tests with aggression)* | *Mean duration of grooming (min), (tests with grooming)* | *Mean duration of body contact (min), (tests with body contact)* |
| 3 | 8 | 5.375 (N=5) | 0.019 (N=1) | 0.054 (N=2) | 3.451 (N=8) |
| 4 | 8 | 1.625 (N=2) | 0.026 (N=2) | 0.014 (N=1) | 4.494 (N=5) |
| 5 | 12 | 3.167 (N=6) | 0.003 (N=1) | 0.077 (N=4) | 3.943 (N=8) |
| 6 | 5 | 3.6 (N=5) | 0 (N=0) | 0.087 (N=1) | 5.771 (N=5) |
| 7 | 4 | 8.5 (N=1) | 0 (N=0) | 0 (N=0) | 1.576 (N=3) |
| 18 | 1 | 3 (N=1) | 0 (N=0) | 0.092 (N=1) | 5.354 (N=1) |
| 20 | 1 | 0 (N=0) | 0 (N=0) | 0 (N=0) | 0.496 (N=1) |
| Means (Totals) | 39 | 3.61 (N=20) | 0.007 (N=4) | 0.046 (N=9) | 3.584 (N=31) |
|  | STDEV | 2.725 | 0.011 | 0.041 | 1.935 |

**Model selection**

**Table S2**: AIC-based model selection results examining reparameterization of age of offspring vs whether offspring had dispersed or not.

| **body contact** | **K** | **-2logL** | **AIC** | **ΔAIC** | **ωAIC** |
| --- | --- | --- | --- | --- | --- |
| dispersed | 6 | -232.99 | 479.16 | 0 | 1 |
| age | 17 | -226.42 | 496.88 | 17.73 | 0 |
| **fight** | **K** | **-2logL** | **AIC** | **ΔAIC** | **ωAIC** |
| dispersed | 6 | 167.41 | -321.66 | 0 | 1 |
| age | 17 | 169.65 | -295.28 | 26.38 | 0 |
| **sniff** | **K** | **-2logL** | **AIC** | **ΔAIC** | **ωAIC** |
| dispersed | 5 | -236.93 | 484.68 | 0 | 1 |
| age | 16 | -231.41 | 503.59 | 18.91 | 0 |
| **groom** | **K** | **-2logL** | **AIC** | **ΔAIC** | **ωAIC** |
| dispersed | 6 | -25.2 | 63.57 | 0 | 1 |
| age | 17 | -31.35 | 106.73 | 43.15 | 0 |

**Table S3. Sniffing:** Results of the Anova models to test whether dispersal and the season have an impact on the frequency of sniffing events in the bush Karoo rat.

*Sniff = season + dispersed*

| *Predictors* | *df* | *Sum sqs* | F value | *p* |
| --- | --- | --- | --- | --- |
| Season | 1 | 18 | 0.723 | 0.398 |
| Dispersed | 1 | 49.2 | 1.978 | 0.164 |
| Season : Dispersed | 1 | 20.5 | 0.822 | 0.368 |

**Table S4. Fighting**: Results of the Anova model to test whether dispersal and the season have an impact on the time spent fighting in the bush Karoo rat.

*Fighting = season + dispersed*

| *Predictors* | *df* | *Sum sqs* | F value | *p* |
| --- | --- | --- | --- | --- |
| Season | 1 | 0.00061 | 0.678 | 0.41 |
| Dispersed | 1 | 0.0015 | 1.656 | 0.2 |
| Weight difference | 1 | 0.00014 | 0.15 | 0.7 |

**Table S5 Grooming:** Results of the Anova models to test whether dispersal and the season have an impact on the time spent grooming in the bush Karoo rat. Factors in bold type indicate significant predictors.

*Grooming = season + dispersed*

| *Predictors* | *df* | *Sum sqs* | F value | *p* |
| --- | --- | --- | --- | --- |
| Season | 1 | 0.00459 | 0.054 | 0.82 |
| Dispersed | 1 | 0.1487 | 1.754 | 0.19 |
| Weight difference | 1 | 0.4105 | 4.841 | **0.03** |

**Table S6. Body contact:** Results of the Anova models to test whether dispersal and the season have an impact on the time spent in body contact in the bush Karoo rat.

*Body contact = season + dispersed*

| *Predictors* | *df* | *Sum sqs* | F value | *p* |
| --- | --- | --- | --- | --- |
| Season | 1 | 0.0236 | 0.0014 | 0.97 |
| Dispersed | 1 | 7.647 | 0.4487 | 0.51 |
| Weight difference | 1 | 24.6384 | 1.4457 | 0.23 |

**Table S7. Neighbor encounter tests - Fighting:** Results of the linear mixed effects models to identify which factors influenced time spent in amicable and aggressive behaviours by bush Karoo rats during dyadic encounter tests with neighbours.

*Fight = season + relatedness + body size difference + ID-focal (random).*

| *Predictors* | *Estimates* | *SE* | 95% CI | *p* |
| --- | --- | --- | --- | --- |
| (Intercept) | -0.007 | 0.018 | -0.043 – 0.029 | 0.692 |
| Season (non-breeding) | 0.016 | 0.013 | -0.0098 – 0.042 | 0.220 |
| Relatedness (non-kin) | 0.009 | 0.015 | -0.02 – -0.04 | 0.551 |
| Weight difference | 0.0003 | 0.001 | -0009 – -0.001 | 0.641 |
| Random effects  *σ*2/ICC | 0.000024/0.009 |  |  |  |
| Marginal  *R*2/conditional *R*2 | 0.03/0.039 |  |  |  |

*σ2, variance of the random effect “ID”; ICC, intraclass coefficient of variation.*

*The marginal R2 considers only the variance of the fixed effects, while the conditional R2 takes both the fixed and random effects into account.*

**Table S8. Neighbor encounter tests – Body contact:** Results of the linear mixed effects models to identify which factors influenced time spent in amicable and aggressive behaviours by bush Karoo rats during dyadic encounter tests with neighbours. Factors in bold type indicate significant predictors.

*Body contact = season + relatedness + body size difference + ID-focal (random).*

| *Predictors* | *Estimates* | *SE* | 95% CI | *p* |
| --- | --- | --- | --- | --- |
| (Intercept) | 4.27 | 1.496 | 1.26 – 7.29 | < **0.01** |
| Season (non-breeding) | 0.57 | 1.094 | -1.63 – 2.80 | 0.60 |
| Relatedness (non-kin) | -2.36 | 1.316 | -5.04 – 0.31 | 0.08 |
| Weight difference | -0.02 | 0.048 | -0.12 – 0.08 | 0.69 |
| Random effects  *σ*2/ICC | 5.078/0.26 |  |  |  |
| Marginal  *R*2/conditional *R*2 | 0.06/0.305 |  |  |  |

*σ2, variance of the random effect “ID”; ICC, intraclass coefficient of variation.*

*The marginal R2 considers only the variance of the fixed effects, while the conditional R2 takes both the fixed and random effects into account.*

**Table S9. Neighbor encounter tests - Grooming:** Results of the linear mixed effects models to identify which factors influenced time spent in amicable and aggressive behaviours by bush Karoo rats during dyadic encounter tests with neighbours.

*Grooms = season + relatedness + body size difference + ID-focal (random).*

| *Predictors* | *Estimates* | *SE* | 95% CI | *p* |
| --- | --- | --- | --- | --- |
| (Intercept) | 0.04 | 0.026 | -0.0097 – 0.095 | 0.09 |
| Season (non-breeding) | 0.008 | 0.019 | -0.0295 – 0.045 | 0.673 |
| Relatedness (non-kin) | -0.02 | 0.022 | -0.0645 – 0.025 | 0.356 |
| Weight difference | -0.0002 | 0.001 | -0.0019 – 0.002 | 0.793 |
| Random effects  *σ*2/ICC | 0.000049/0.009 |  |  |  |
| Marginal  *R*2/conditional *R*2 | 0.02/0.029 |  |  |  |

*σ2, variance of the random effect “ID”; ICC, intraclass coefficient of variation.*

*The marginal R2 considers only the variance of the fixed effects, while the conditional R2 takes both the fixed and random effects into account.*

**Table S10. Sniffing events during encounter tests:** Results of the generalized linear mixed effects models to identify which factors influenced the frequency of sniffing events exhibited by bush Karoo rats during dyadic encounter tests towards neighbours. Factors in bold type indicate significant predictors.

*Sniff = season * relatedness + body size difference + ID-focal (random) + ID-stimulus(random).*

| *Predictors* | *Estimates* | *SE* | 95% CI | *p* |
| --- | --- | --- | --- | --- |
| (Intercept) | 0.42 | 0.35 | -0.29 – 1.08 | 0.22 |
| Season (non-breeding) | 0.73 | 0.27 | 0.22 – 1.29 | **< 0.01** |
| Relatedness (non-kin) | 0.54 | 0.36 | -0.17 – 1.27 | 0.14 |
| Weight difference | -0.02 | 0.008 | -0.04 - -0.004 | **0.02** |
| Season (non-breeding) : Relatedness (non-kin) | -0.63 | 0.43 | -1.49 – 0.21 | 0.14 |
| Random effects  *σ*2/ICC | 2.10/0.83 |  |  |  |
| Marginal  *R*2/conditional *R*2 | 0.061/0.839 |  |  |  |

*σ2, variance of the random effect “ID”; ICC, intraclass coefficient of variation.*

*The marginal R2 considers only the variance of the fixed effects, while the conditional R2 takes both the fixed and random effects into account.*

**Table S11. Latency to first aggression during intruder tests:** Results of the linear mixed effects models to identify which factors influenced the latency to the first aggressive behaviour with a stimulus during intruder tests towards neighbours.

*Latency to first aggression = season * relatedness + body size difference + ID-focal (random).*

| *Predictors* | *Estimates* | *SE* | 95% CI | *p* |
| --- | --- | --- | --- | --- |
| (Intercept) | 4.286 | 1.69 | 0.08 – 7.67 | **< 0.01** |
| Season (non-breeding) | 4.287 | 1.82 | 0.61 – 7.95 | **0.02** |
| Relatedness (non-kin) | -3.06 | 1.64 | -6.47 – 0.24 | 0.07 |
| Weight difference | 0.05 | 0.03 | -0.0091 – 0.118 | 0.08 |
| Season (non-breeding) : Relatedness (non-kin) | 1.52 |  | -2.59 – 5.65 | 0.46 |
| Random effects  *σ*2/ICC | 11.07 /0.52 |  |  |  |
| Marginal  *R*2/conditional *R*2 | 0.348/0.688 |  |  |  |

*σ2, variance of the random effect “ID”; ICC, intraclass coefficient of variation.*

*The marginal R2 considers only the variance of the fixed effects, while the conditional R2 takes both the fixed and random effects into account.*

**Table S12. Latency to approach cage during intruder tests:** Results of the linear mixed effects models to identify which factors influenced the latency to approach a cage with a stimulus during intruder tests towards neighbours.

*Latency to approach = season * relatedness + body size difference + ID-focal (random).*

| *Predictors* | *Estimates* | *SE* | 95% CI | *p* |
| --- | --- | --- | --- | --- |
| (Intercept) | 0.89 | 1.70 | -2.68 – 4.32 | 0.60 |
| Season (non-breeding) | 2.88 | 1.82 | -0.94 – 6.69 | 0.12 |
| Relatedness (non-kin) | -0.98 | 1.61 | -4.27 – 2.29 | 0.55 |
| Weight difference | 0.04 | 0.03 | -0.02 – 0.01 | 0.17 |
| Season (non-breeding) : Relatedness (non-kin) | 3.02 | 1.97 | -1.01 – 7.06 | 0.14 |
| Random effects  *σ*2/ICC | 13.22/0.556 |  |  |  |
| Marginal  *R*2/conditional *R*2 | 0.236/0.661 |  |  |  |

*σ2, variance of the random effect “ID”; ICC, intraclass coefficient of variation.*

*The marginal R2 considers only the variance of the fixed effects, while the conditional R2 takes both the fixed and random effects into account.*

**Table S13. Charging events during intruder tests:** Results of the generalized linear mixed effects models to identify which factors influenced the frequency of charging events exhibited by bush Karoo rats during intruder tests towards neighbours. Factors in bold type indicate significant predictors.

*Charging = season * relatedness + body size difference + ID-focal (random).*

| *Predictors* | *Estimates* | *SE* | 95% CI | *p* |
| --- | --- | --- | --- | --- |
| (Intercept) | 1.55 | 0.28 | 0.97 – 2.16 | **< 0.001** |
| Season (non-breeding) | -1.92 | 0.44 | -2.899 - -1.10 | < **0.001** |
| Relatedness (non-kin) | 0.75 | 0.22 | 0.32 – 1.20 | < **0.001** |
| Weight difference | -0.0056 | 0.0049 | -0.016 – 0.004 | 0.26 |
| Season (non-breeding) : Relatedness (non-kin) | 0.54 | 0.39 | -0.21 – 1.37 | 0.17 |
| Random effects  *σ*2/ICC | 0.41/0.604 |  |  |  |
| Marginal  *R*2/conditional *R*2 | 0.620/0.849 |  |  |  |

*σ2, variance of the random effect “ID”; ICC, intraclass coefficient of variation.*

*The marginal R2 considers only the variance of the fixed effects, while the conditional R2 takes both the fixed and random effects into account.*

**Table S14. Number of trill sounds during intruder tests:** Results of the linear mixed effects models to identify which factors influenced the number of trill sounds with a stimulus during intruder tests towards neighbours.

*Trills = season * relatedness + body size difference + ID-focal (random).*

| *Predictors* | *Estimates* | *SE* | 95% CI | *p* |
| --- | --- | --- | --- | --- |
| (Intercept) | 21.997 | 14.1 | -4.73 – 48.83 | 0.13 |
| Season (non-breeding) | -9.66 | 16.94 | -41.62 – 22.76 | 0.57 |
| Relatedness (non-kin) | 9.53 | 18.2 | -25.45 – 44.77 | 0.61 |
| Weight difference | -0.38 | 0.27 | -0.90 – 0.13 | 0.17 |
| Season (non-breeding) : Relatedness (non-kin) | 7.48 | 23.25 | -37.60 – 52.53 | 0.75 |
| Random effects  *σ*2/ICC | 101.6/0.65 |  |  |  |
| Marginal  *R*2/conditional *R*2 | 0.121/0.178 |  |  |  |

*σ2, variance of the random effect “ID”; ICC, intraclass coefficient of variation.*

*The marginal R2 considers only the variance of the fixed effects, while the conditional R2 takes both the fixed and random effects into account.*

**Table S15. Time spent at a cage during intruder tests:** Results of the linear mixed effects models to identify which factors influenced the time spent at a cage of the stimulus during intruder tests. Factors in bold type indicate significant predictors.

*Time at cage = season * relatedness + body size difference + ID-focal (random).*

| *Predictors* | *Estimates* | *SE* | 95% CI | *p* |
| --- | --- | --- | --- | --- |
| (Intercept) | 5.08 | 1.25 | 2.54 – 7.59 | **< 0.001** |
| Season (non-breeding) | -2.11 | 1.42 | -4.95 – 0.75 | 0.14 |
| Relatedness (non-kin) | 4.63 | 1.37 | 1.76 – 7.39 | **< 0.01** |
| Weight difference | -0.03 | 0.02 | -0.07 – 0.024 | 0.30 |
| Season (non-breeding) : Relatedness (non-kin) | -1.93 | 1.71 | -5.43 – 1.59 | 0.27 |
| Random effects  *σ*2/ICC | 4.38/0.354 |  |  |  |
| Marginal  *R*2/conditional *R*2 | 0.381/0.600 |  |  |  |

*σ2, variance of the random effect “ID”; ICC, intraclass coefficient of variation.*

*The marginal R2 considers only the variance of the fixed effects, while the conditional R2 takes both the fixed and random effects into account.*

**Supplementary figures**

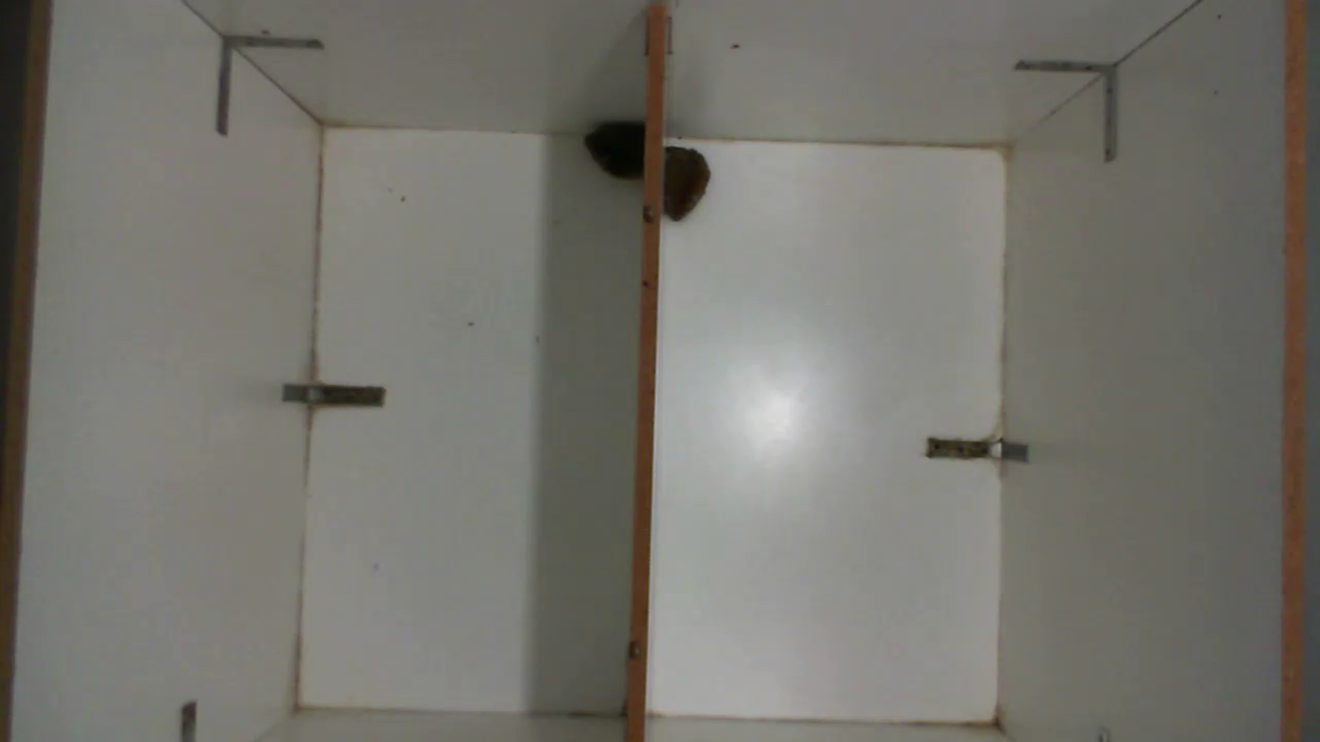

**Figure S 1:** Picture showing the dyadic encounter test neutral arena with a partition in between. In the picture, two rats are seen sitting beside the partition.

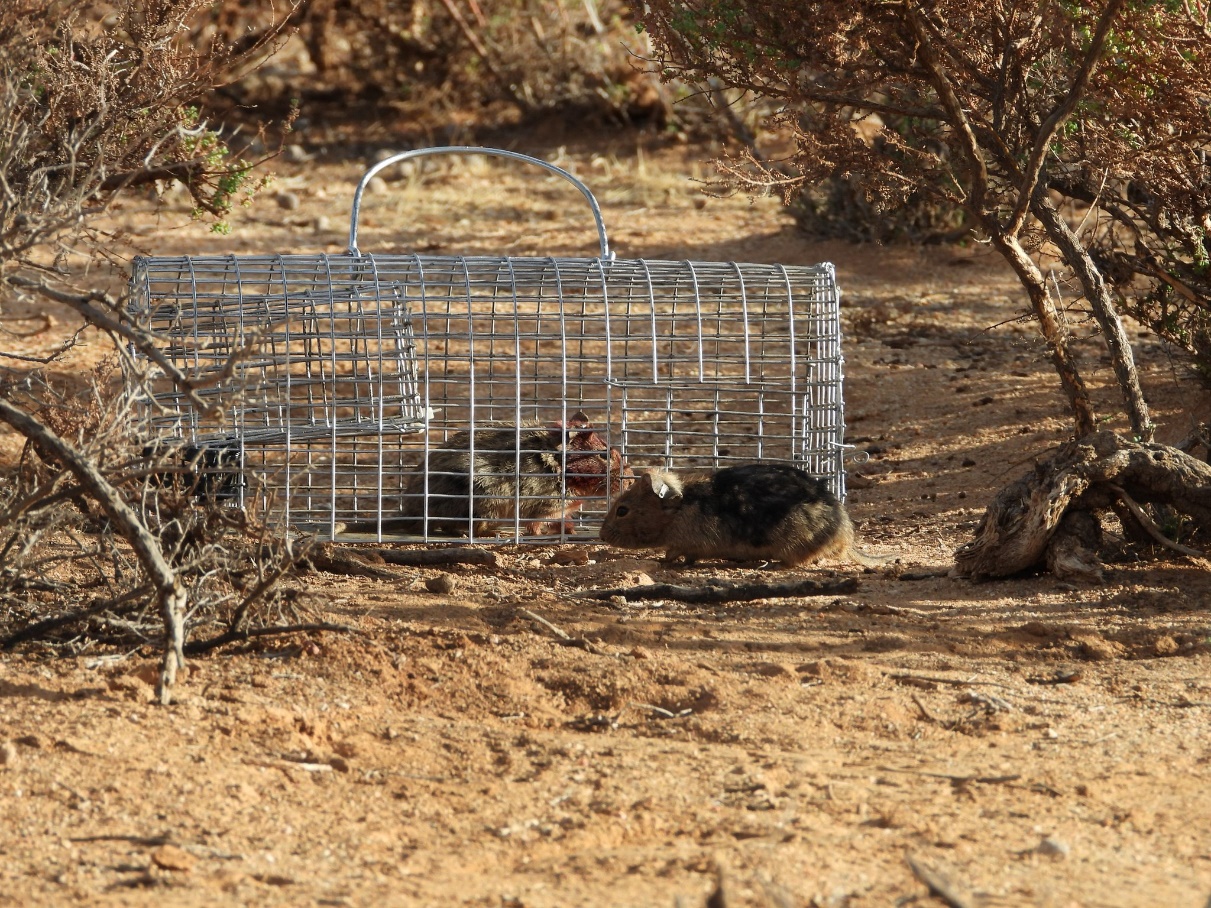

**Figure S 2:** Figure showing the set-up for the field intruder tests. The focal rat is seen outside of the cage and the stimulus is inside the cage.

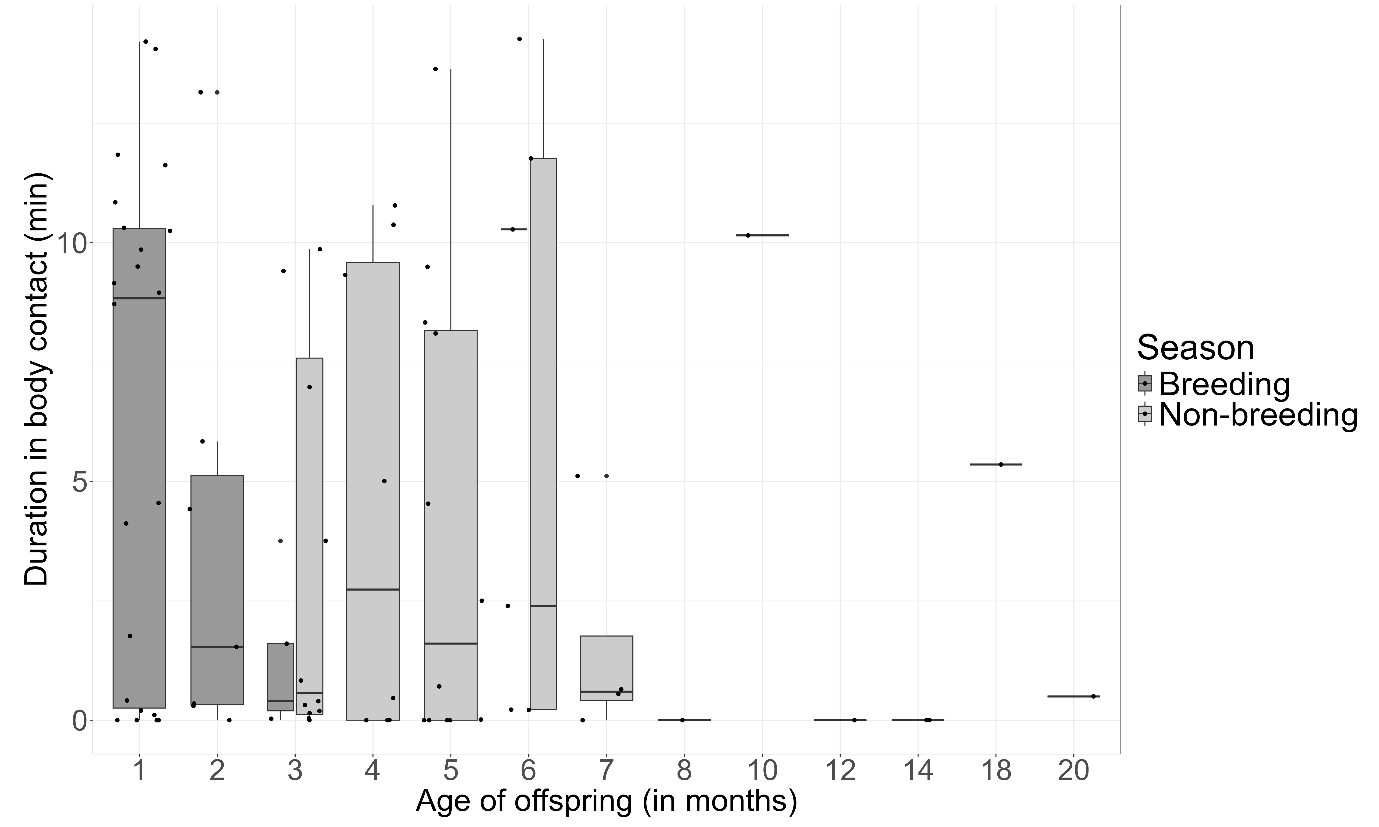

**Figure S 3**: The duration spent in body contact during 15min observations by female bush Karoo rats according to the age of the offspring, for both seasons. Boxplots show median and 1st and 3rd quartiles, the whiskers represent the minimum and maximum of the outlier data and points represent individual values (breeding: n = 40, non-breeding: n = 39).

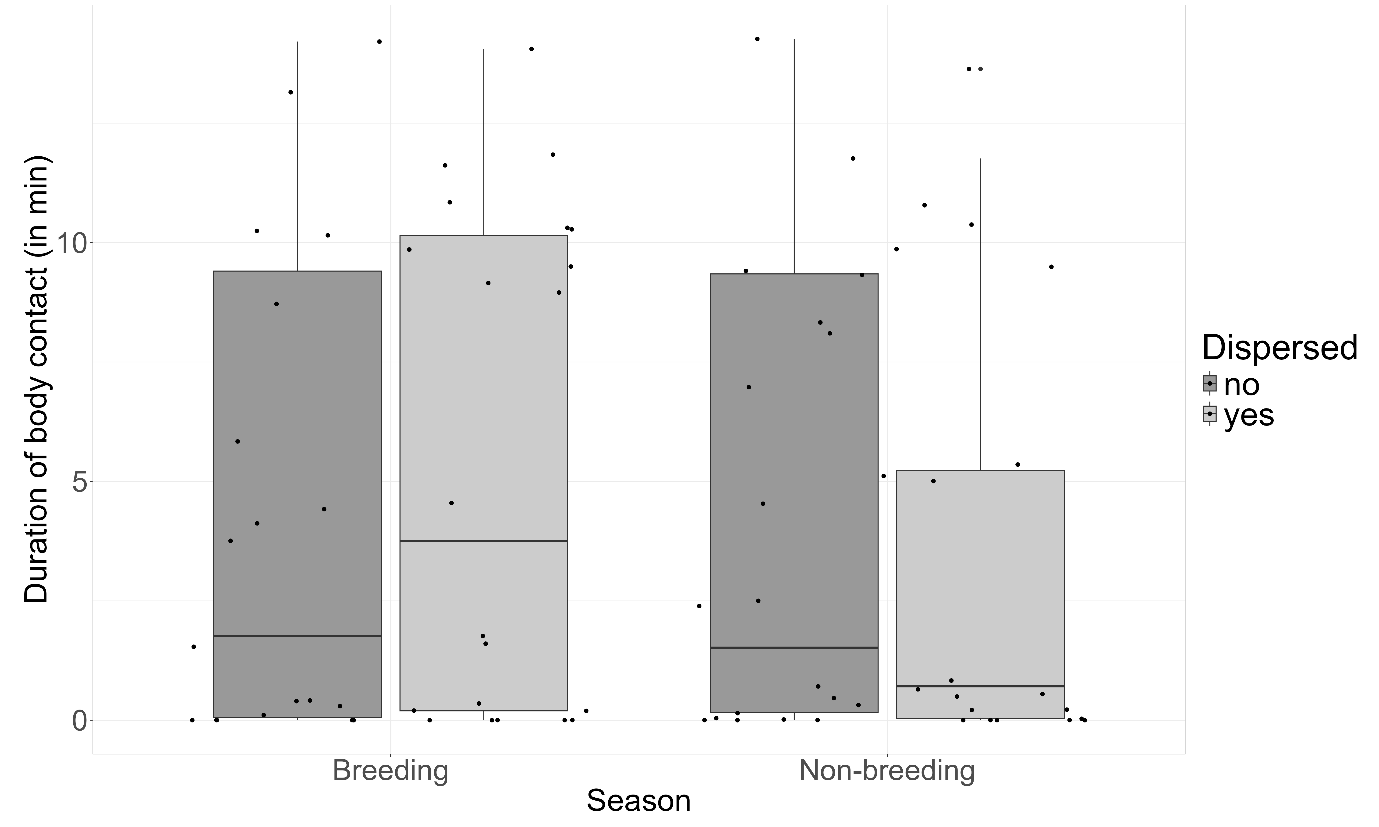

**Figure S 4**: The duration spent in body contact by female bush Karoo rats according to whether offspring had dispersed or not, for both seasons. Boxplots show median and 1st and 3rd quartiles, the whiskers represent the minimum and maximum of the outlier data and points represent individual values (breeding: n = 40, non-breeding: n = 39).

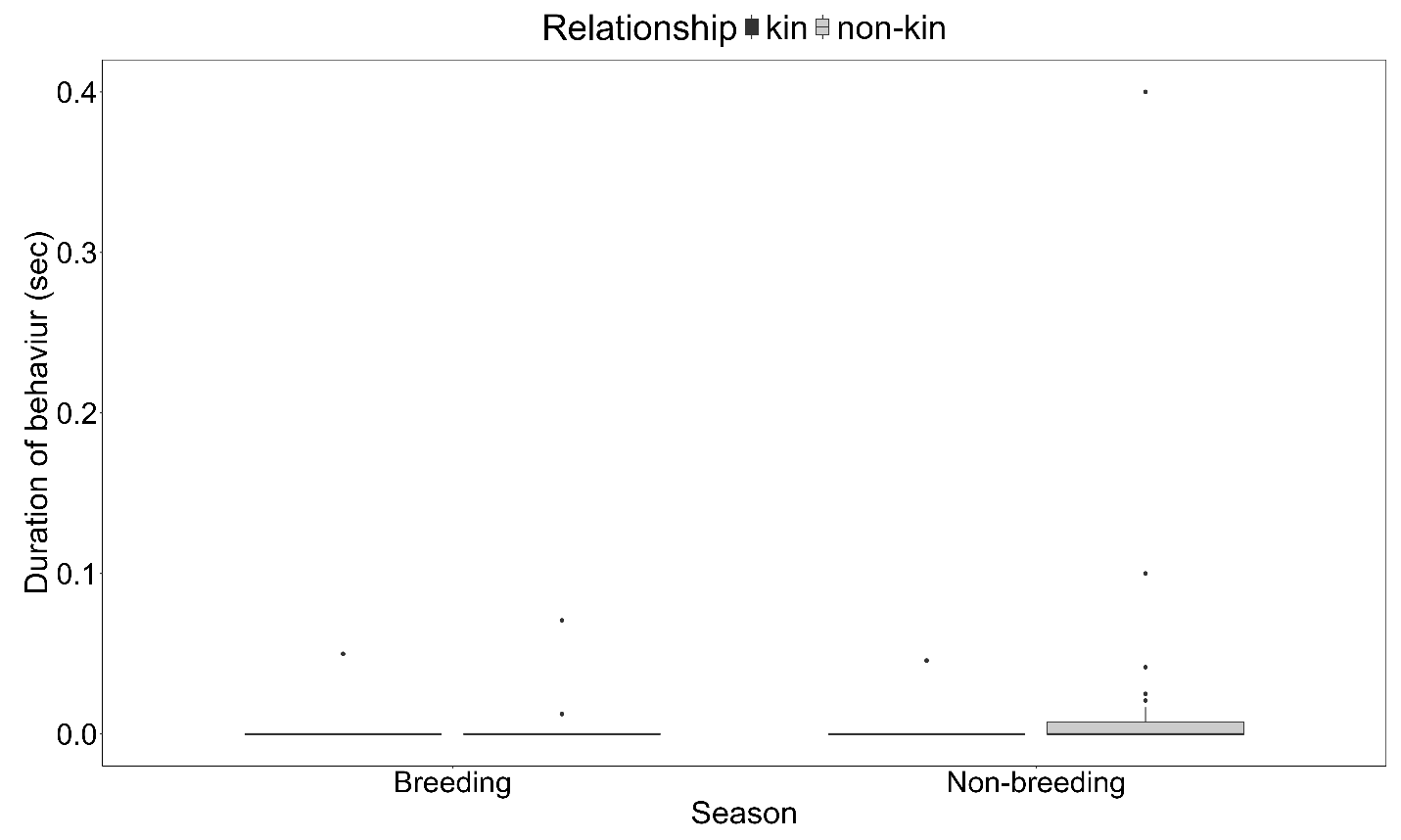

**Figure S 5**: The duration spent fighting by female bush Karoo rats during dyadic encounter tests for both seasons. Boxplots show median and 1st and 3rd quartiles, the whiskers represent the minimum and maximum of the outlier data and points represent individual values (breeding: n = 40, non-breeding: n = 39).

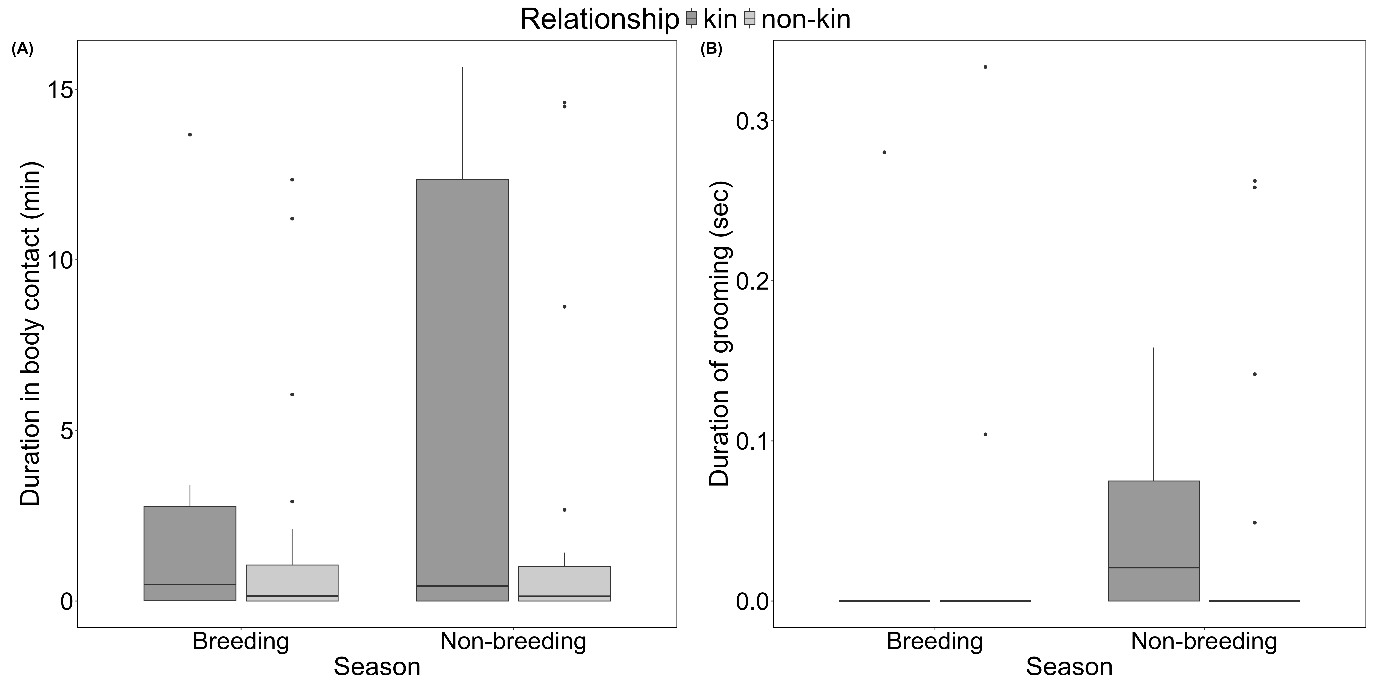

**Figure S 6**: The duration of body contact (A) and grooming (B) behaviours between neighbouring female bush Karoo rats during dyadic encounter tests. Boxplots show median and 1st and 3rd quartiles, the whiskers represent the minimum and maximum of the outlier data and points represent individual values.

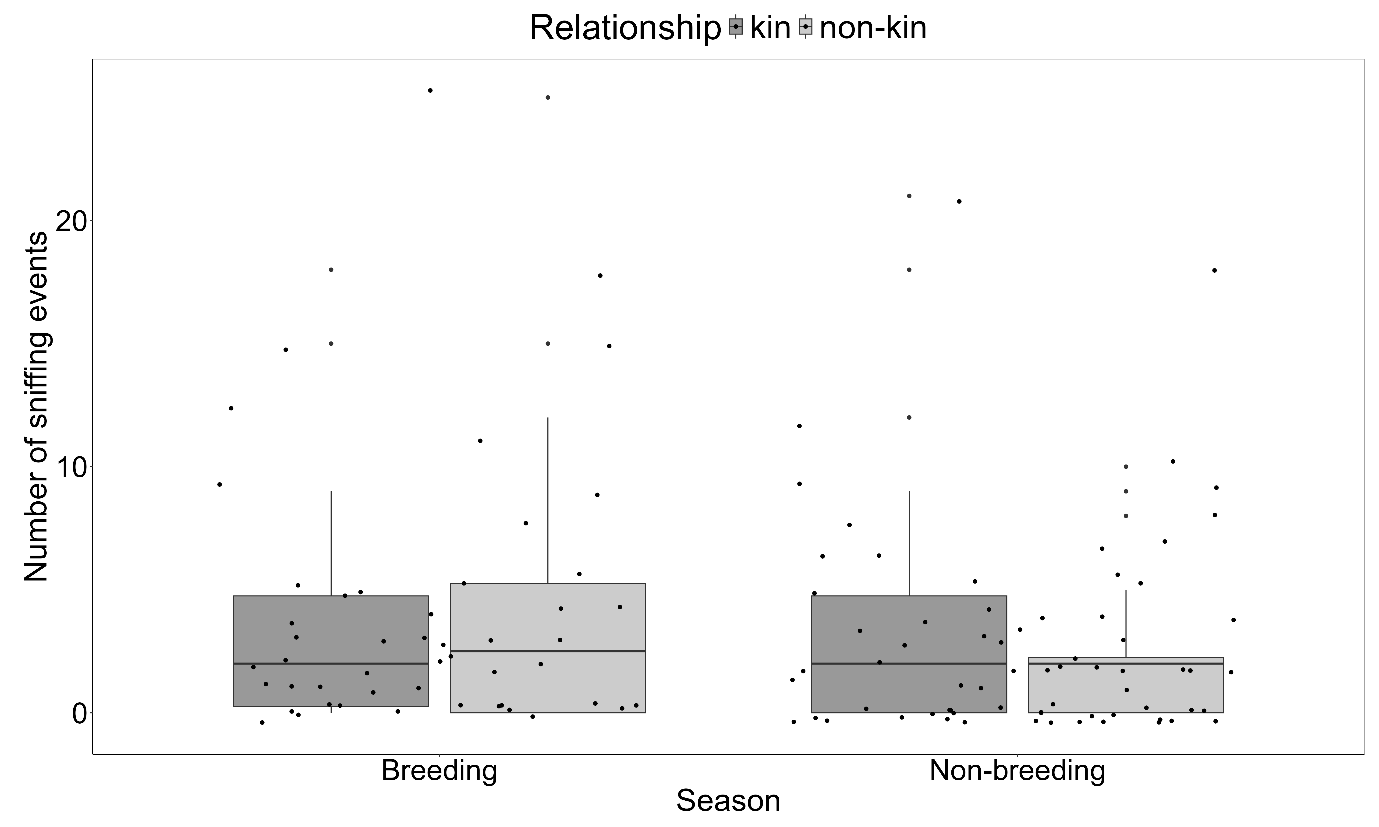

**Figure S 7**: The number of sniffing events between neighbouring female bush Karoo rats during dyadic encounter tests. Boxplots show median and 1st and 3rd quartiles, the whiskers represent the minimum and maximum of the outlier data and points represent individual values.

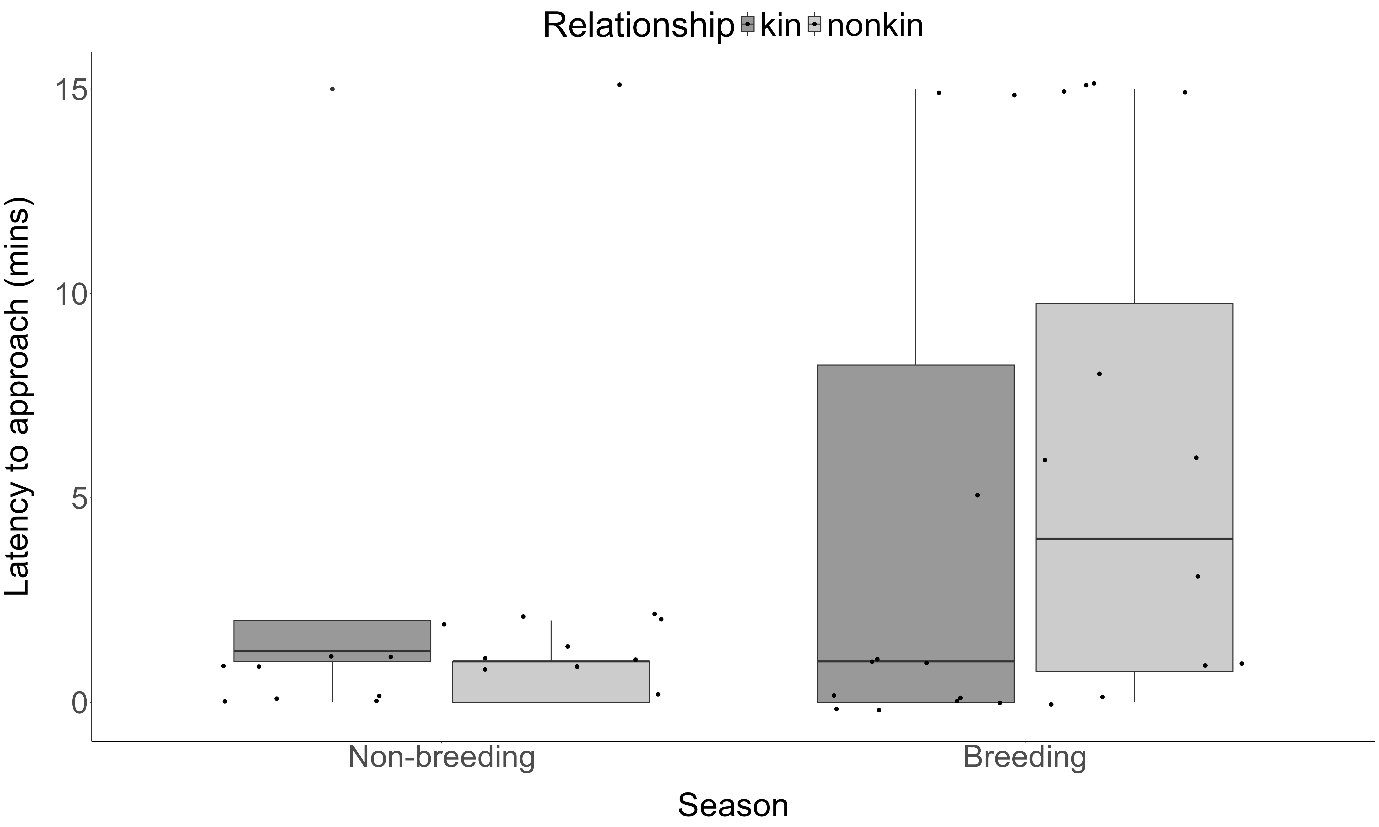

**Figure S 8:** A measure of the latency to approach a cage during intruder tests based on the relationship between individuals. Boxplots show median and 1st and 3rd quartiles, the whiskers represent the minimum and maximum of the outlier data and points represent individual values.

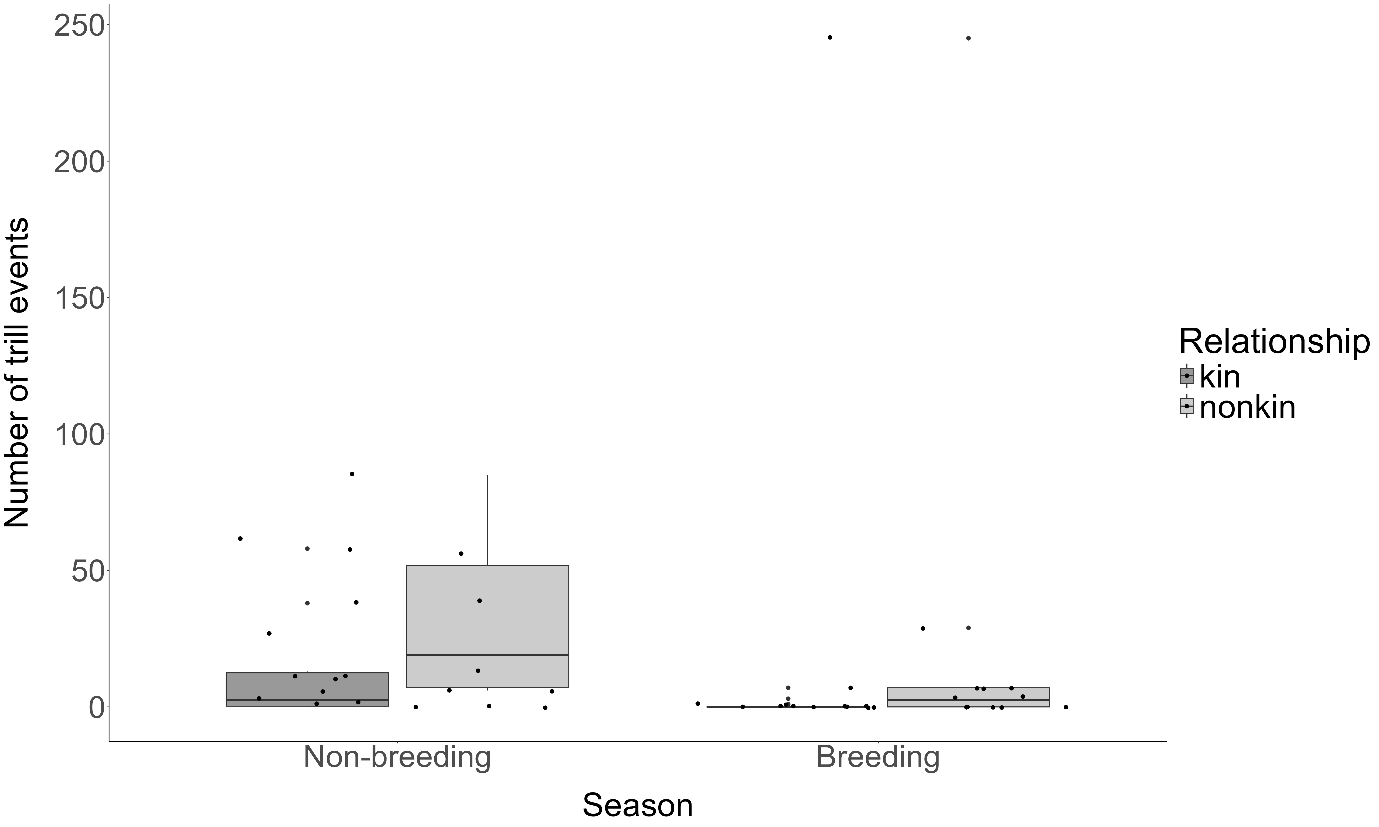

**Figure S 9:** The number of trilling events measured during intruder tests. Boxplots show median and 1st and 3rd quartiles, the whiskers represent the minimum and maximum of the outlier data and points represent individual values.

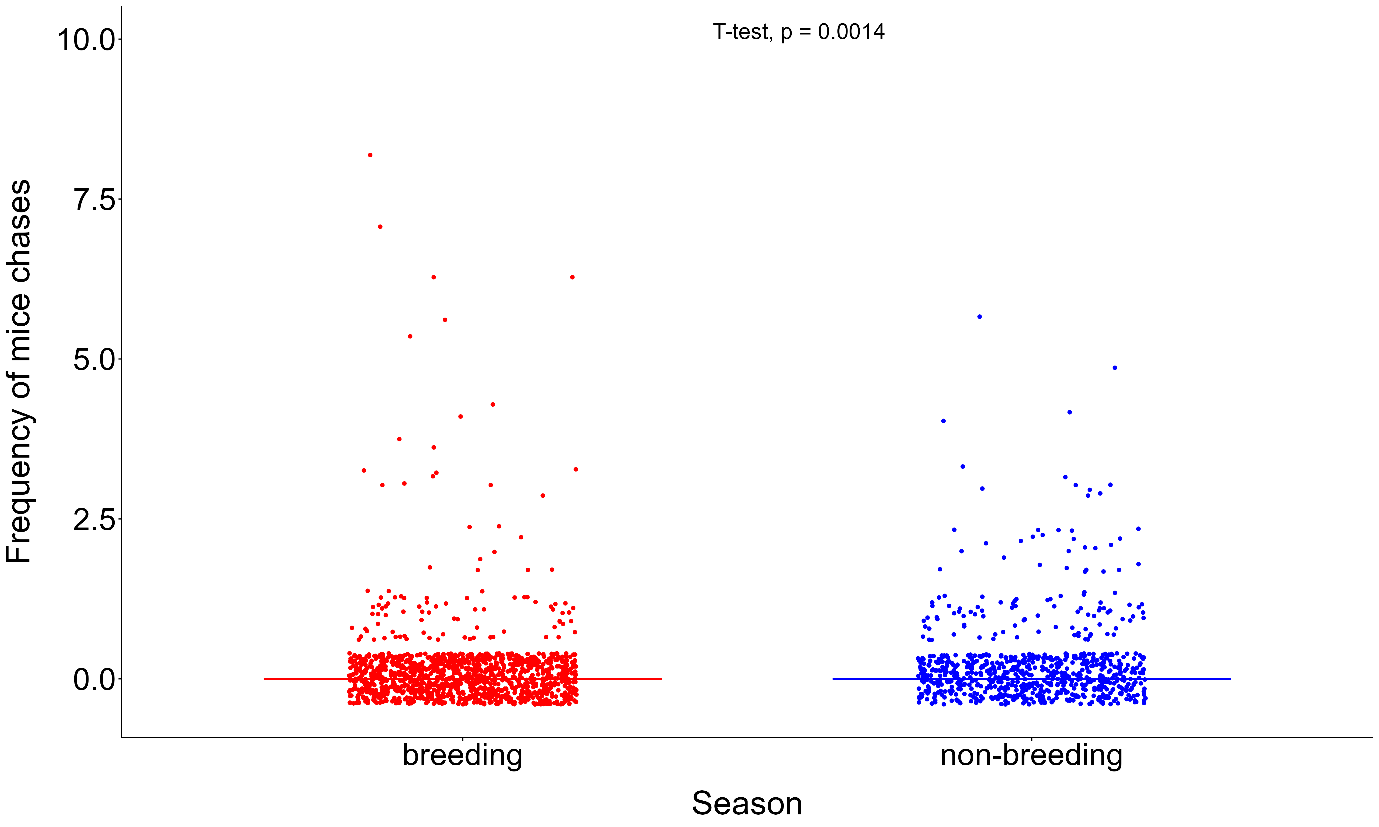

**Figure S 10:** The frequency of chasing events against striped mice.

**Additional tables on models run using age of offspring as a variable.**

**Table S3.1. Sniffing:** Results of the Anova models to test whether age of offspring and the season have an impact on the frequency of sniffing events in the bush Karoo rat.

*Sniff = season + age of offspring*

| *Predictors* | *df* | F value | *p* |
| --- | --- | --- | --- |
| Season | 1 | 0.711 | 0.402 |
| Age offspring (months) | 12 | 1.044 | 0.422 |
| Season : Age offspring (months) | 1 | 0.001 | 0.978 |

**Table S4.1. Fighting**: Results of the Anova model to test whether age of offspring and the season have an impact on the time spent fighting in the bush Karoo rat.

*Fighting = season + age of offspring*

| *Predictors* | *df* | F value | *p* |
| --- | --- | --- | --- |
| Season | 1 | 0.619 | 0.438 |
| Age offspring (months) | 12 | 0.418 | 0.953 |
| Weight difference | 1 | 1.101 | 0.298 |

**Table S5.1 Grooming:** Results of the Anova models to test whether age of offspring and the season have an impact

on the time spent grooming in the bush Karoo rat.

*Grooming = season + age of offspring*

| *Predictors* | *df* | F value | *p* |
| --- | --- | --- | --- |
| Season | 1 | 0.029 | 0.865 |
| Age offspring (months) | 12 | 0.105 | 0.9999 |
| Weight difference | 1 | 0.732 | 0.396 |

**Table S6.1. Body contact:** Results of the Anova models to test whether age of offspring and the season have an impact on the time spent in body contact in the bush Karoo rat.

*Body contact = season + age of offspring*

| *Predictors* | *df* | F value | *p* |
| --- | --- | --- | --- |
| Season | 1 | 0.171 | 0.68 |
| Age offspring (months) | 12 | 1.244 | 0.27 |
| Weight difference | 1 | 0.138 | 0.71 |
